# Segmenting surface boundaries using luminance cues: Underlying mechanisms

**DOI:** 10.1101/2020.06.27.175505

**Authors:** Christopher DiMattina, Curtis L. Baker

## Abstract

Segmenting scenes into distinct surfaces is a basic visual perception task, and luminance differences between adjacent surfaces often provide an important segmentation cue. However, mean luminance differences between two surfaces may exist without any sharp change in albedo at their boundary, but rather from differences in the proportion of small light and dark areas within each surface, e.g. texture elements, which we refer to as a *luminance texture boundary*. Here we investigate the performance of human observers segmenting luminance texture boundaries. We demonstrate that a simple model involving a single stage of filtering cannot explain observer performance, unless it incorporates contrast normalization. Performing additional experiments in which observers segment luminance texture boundaries while ignoring super-imposed luminance step boundaries, we demonstrate that the one-stage model, even with contrast normalization, cannot explain performance. We then present a Filter-Rectify-Filter (FRF) model positing two cascaded stages of filtering, which fits our data well, and explains observers’ ability to segment luminance texture boundary stimuli in the presence of interfering luminance step boundaries. We propose that such computations may be useful for boundary segmentation in natural scenes, where shadows often give rise to luminance step edges which do not correspond to surface boundaries.

## INTRODUCTION

Detecting boundaries separating distinct surfaces is a crucial first step for segmenting the visual scene into regions. Since different surfaces generally reflect different proportions of the illuminant, luminance differences provide a highly informative cue for boundary detection in natural images (Mely, Kim, McGill, Guo, & Serre, 2016; DiMattina, Fox, & Lewicki, 2012; Martin, Fowlkes, & Malik, 2004; Marr, 1982). Inspired by physiological findings (Hubel & Wiesel, 1962; Parker & Hawken, 1988), a commonly assumed computational model of luminance boundary detection is a Gabor-shaped linear spatial filter of appropriate spatial scale and orientation (or a multi-scale population of filters) detecting a localized change in luminance near the boundary (**Fig. 1a, b**) (Elder & Sachs, 2004; Marr, 1982). However, in many natural scenes, two distinct surfaces may visibly differ in their mean regional luminance without giving rise to any sharp change in luminance at their boundary. This situation is illustrated in **Fig. 1d**, which shows two juxtaposed textures from the Brodatz database (Brodatz, 1966). Clearly, a large-scale Gabor filter defined on the scale of the whole image as in **Fig. 1a** can certainly provide some information about a difference in average luminance between the two surfaces. However, it is unknown whether other mechanisms may be better suited to detect regional luminance differences at such boundaries.

**Figure 1:**
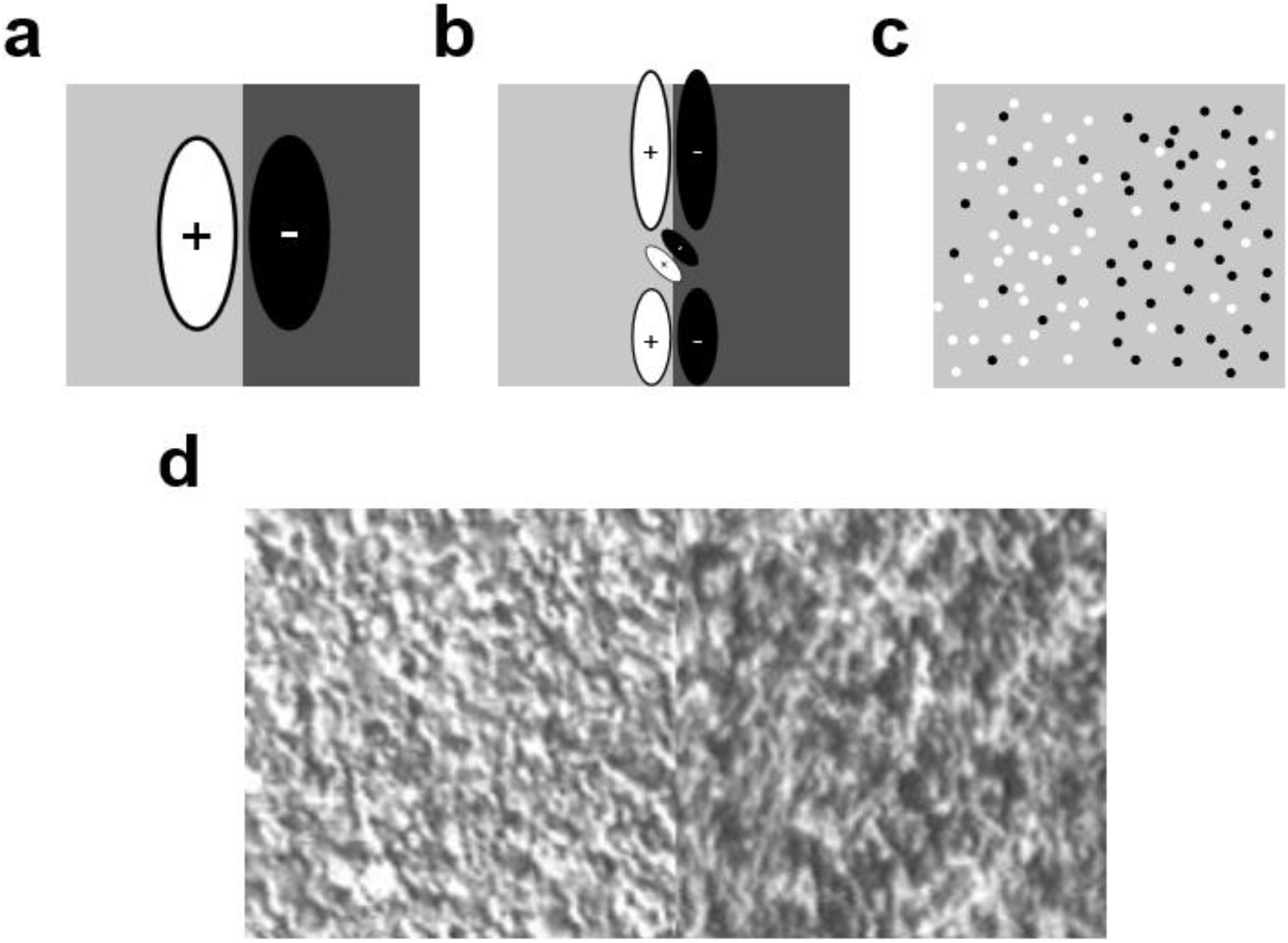
Boundaries without luminance step edges. (**a**) A luminance step boundary (LSB) and a simple detection model in which a linear Gabor filter measures the regional luminance difference. (**b**) Model similar to that in (a) where the LSB is analyzed by multiple Gabor filters at varying spatial scales. (**c**) Example of luminance texture boundary (LTB). The luminance difference is defined by differing proportions of black and white micropatterns on each side of the boundary, with no sharp luminance change at the boundary. (**d**) Two juxtaposed textures from the Brodatz database. Although there is clearly a regional difference in luminance, there is no sharp luminance change at the boundary.

In order to address this question, we propose a basic taxonomy of two different ways that luminance cues can define region boundaries. *Luminance step boundaries* (LSBs) are defined by uniform regional differences in luminance, as in **Fig. 1a**. *Luminance texture boundaries* (LTBs) are defined by differing proportions of dark and light texture elements or micro-patterns on two adjacent surfaces (**Fig. 1c**). Note that for the artificial LTB shown in **Fig. 1c** there are no textural cues present other than the proportions of dark and light elements on each side of the boundary. Given that regional luminance differences can arise from either LSBs or LTBs, it is of interest to understand whether or not similar mechanisms are employed when segmenting these boundaries, and how LTBs and LSBs interact when both are present, as for example when a cast shadow falls upon a scene region containing one or more surface boundaries.

A number of studies have investigated detection of “first-order” luminance step boundaries (Elder & Sachs, 2004; McIlhagga & May, 2012; McIlhagga, 2018; McIlhagga & Mullen, 2018), as well as detection and segmentation of “second-order” texture boundaries having no luminance difference but differences in texture contrast (Dakin & Mareschal, 2000; DiMattina & Baker, 2019), density (Zavitz & Baker, 2014), orientation (Wolfson & Landy, 1995), polarity (Motoyoshi & Kingdom, 2007) or phase (Hansen & Hess, 2006). However, the segmentation of first-order luminance texture boundaries, and the underlying computations, are poorly understood.

In this study, we characterize perceptual segmentation of LTBs (**Experiment 1**) and demonstrate that simple regional luminance difference computation cannot readily explain their segmentation (**Experiments 2, 3**). We demonstrate the robustness of LTB segmentation to variations in contrast of texture elements, and demonstrate an excellent fit to the data with a psychometric function incorporating divisive contrast normalization (**Experiment 3**). We show that when both cues are present, observers can ignore masking LSBs having orthogonal orientations when segmenting LTBs using proportion of imbalanced patterns as a segmentation cue (**Experiment 4**). However, the presence of a masking LSB having a congruent orientation with the target LTB can in some cases enhance or impair performance (depending on relative phase), suggesting some degree of pre-attentive interaction between cues. An additional experiment further demonstrated the robustness of LTB segmentation to masking LSBs (**Experiment 5**).

We test the ability of a simple model positing a single stage of filtering which fit the data well in **Experiments 2, 3** but it fails to fully explain the results of **Experiment 4**, suggesting that LTBs and LSBs are segmented by distinct underlying mechanisms. We define and fit a “filter-rectify-filter” (FRF) model positing two stages of filtering to data from **Experiment 4**, and show that this model successfully accounts for observer performance in the task. Previous studies of second-order vision have fit psychophysical data with FRF models (DiMattina & Baker, 2019; Zavitz & Baker, 2013, 2014), but here we show that the FRF model can also account for the ability of observers to extract first-order (luminance) information in the presence of masking LSB stimuli. We propose that such mechanisms may be useful for performing boundary segmentation in natural vision, where extraneous stimuli such as shadows often give rise to LSB stimuli which do not correspond to surface boundaries.

## METHODS

### Stimuli

#### Luminance texture boundaries

Luminance texture boundary (LTB) stimuli were created by placing different proportions of non-overlapping black (B) and white (W) micropatterns on opposite halves of a circular disc, with the boundary separating the two regions oriented left-oblique (−45 degrees w.r.t. vertical) or right-oblique (+45 deg. w.r.t. vertical), as shown in **Fig. 2a**. The proportions of black and white micro-patterns on each side of the LTB was parameterized by the proportion *π_U_* of “unbalanced” micro-patterns on each side of the disc (i.e., those not having a counterpart of opposite luminance polarity). Note that *π_U_* can range from 0 (indicating an equal number of black and white micropatterns on both sides) to +1 (opposite colors on opposite sides).

**Figure 2:**
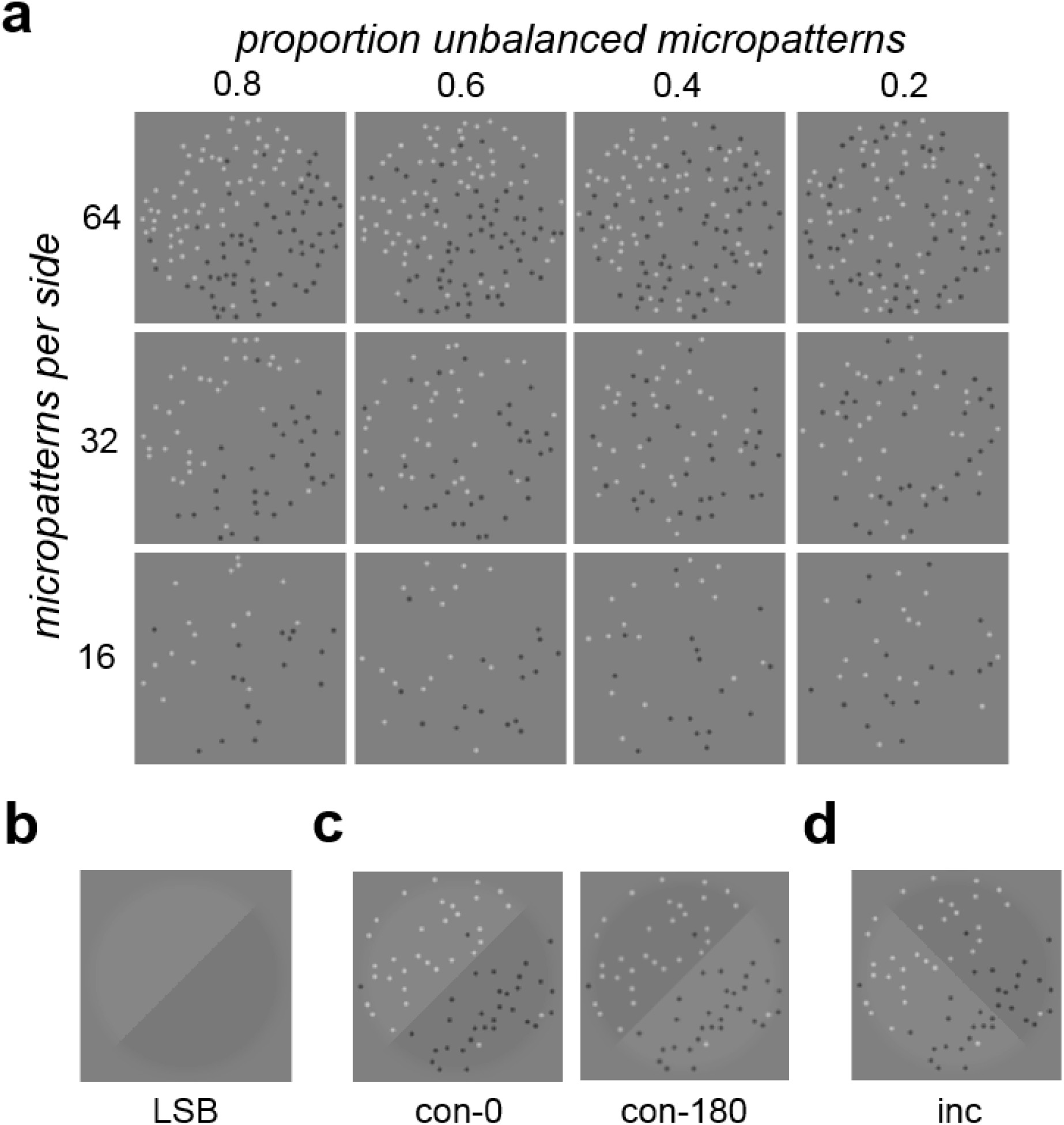
Stimulus images. (**a**) Examples of luminance texture boundary (LTB) stimuli used in this study, shown for varying densities (16, 32, 64 micropatterns on each side of boundary) and proportion unbalanced micropatterns (*π_U_* = 0.2, 0.4, 0.6, 0.8). For all of these example stimulus images, the boundary is right oblique. (**b**) Luminance step boundary (LSB) stimulus. (**c**) Stimulus image examples with LTB and LSB having the same orientation (*congruent*), either phase-aligned (**con-0**) or opposite-phase (**con-180**). (**d**) Example image having superimposed, orthogonal (*incongruent*) luminance texture (right-oblique) and luminance step (left-oblique) boundaries (**inc**).

For the experiments described here, we employed a 256 x 256 pixel stimulus subtending 4 deg. visual angle (dva). An equal number (16, 32 or 64) of non-overlapping micro-patterns were randomly placed on each side of the boundary, with each micro-pattern being an 8 pixel Gaussian (σ = 2 pixels). Unless otherwise specified, the micro-pattern maximum amplitude *A* was set to +/− 0.25 (W/B) dimensionless luminance units with respect to the gray mid-point (0.5), so these micropatterns were clearly visible. Michealson contrast *C*_*M*_ = (*L_max_* − *L_min_*)⁄(*L_max_* + *L_min_*) of the LTB stimuli is related to the maximum micro-pattern amplitude *A* by *C*_*M*_ = 2*A*. In some experiments, we set *A* = +/−0.1 (roughly 3-4 times LTB contrast detection threshold) to create a more difficult task due to reduced visibility of the micro-patterns. Stimuli were designed to have zero luminance difference across the diagonal perpendicular to the region boundary (*anti-diagonal*), so that the only available luminance cue was that across the boundary defining the stimulus. For each stimulus we randomized the modulation *envelope phase* to either ϕ = 0 degrees (left side brighter) or ϕ = 180 degrees (right side brighter).

### Luminance step boundaries

We also characterized performance on our identification task with luminance step boundary (LSB) stimuli, like that shown in **Fig. 2b**. LSB stimuli, produced by multiplying an obliquely oriented step edge by a cosine-tapered circular disc, were also 256 x 256 pixels and scaled to subtend 4 dva. The detectability of this edge was varied by manipulating its Michealson contrast *C*_*M*_, and again envelope phase was randomized.

### Observers

Two groups of observers participated as psychophysical observers in these experiments. The first group consisted of N = 3 observers who were highly experienced with the segmentation tasks. One of these observers was author CJD, and the other two (KNB, ERM) were undergraduate members of the Computational Perception Laboratory who were naïve to the purpose of the experiments. The second group was comprised of N = 17 naïve, inexperienced observers recruited from undergraduate FGCU Psychology classes, as well as N = 1 student from the Computational Perception Lab. All observers had normal or corrected-to-normal visual acuity. All observers gave informed consent, and all experimental procedures were approved by the FGCU IRB (Protocol number 2014-01), in accordance with the Declaration of Helsinki.

### Visual Displays

Stimuli were presented in a dark room on a 1920×1080, 120 Hz gamma-corrected Display++ LCD Monitor (Cambridge Research Systems LTD®) with mid-point luminance of 100 cd/m^2^. This monitor was driven by an NVIDA GeForce® GTX-645 graphics card, and experiments were controlled by a Dell Optiplex® 9020 running custom-authored software written in MATLAB® making use of Psychtoolbox-3 routines (Brainard, 1997; Pelli, 1997). Observers were situated 133 cm from the monitor using a HeadSpot® chin-rest, so that the 256×256 stimuli subtended approximately 4 deg. of visual angle.

### Experimental Protocols

#### Experiment 1: Segmentation thresholds for LTBs and LSBs

Towards the larger goal of determining whether the two kinds of luminance boundaries (LTB) are segmented using the same mechanisms, we started by characterizing observers’ segmentation thresholds for both kinds of stimulus. In this and subsequent experiments, the psychophysical task was a single-interval classification task, in which the observer classifies a single displayed stimulus as belonging to one of two categories (L/R oblique).

To study the effects of the number of unbalanced micro-patterns on segmentation (**Experiment 1a**), luminance texture boundaries with 32 micro-patterns on each side were presented at 9 evenly spaced values of *π_U_* from 0 to 1 in steps of 0.125 - example stimulus images are shown in **Fig. 2a**. Observers performed 250 psychophysical trials starting at the highest level, with the stimulus level being adjusted using a standard 1-up, 2-down staircase procedure, focusing trials near stimulus levels yielding 70.71% correct responses (Leek, 2001). Pilot studies with N = 3 experienced observers (CJD, ERM, KNB) showed similar thresholds for 32 and 64 micro-patterns, and somewhat higher thresholds for 16 micro-patterns (**Supplementary Fig. S1**), justifying the use of 32 micro-patterns as our default micro-pattern density.

Luminance step boundaries (LSBs, **Fig. 2b**) were defined by a luminance step oriented either left- or right-oblique, multiplied by a circular window with cosine tapering (Zavitz & Baker, 2013). LSBs were defined by their Michaelson contrast *C*_*M*_ with respect to the luminance midpoint. LSBs were presented at Michealson contrasts in 11 logarithmic steps from *C*_*M*_= 10^−2.7^ to 10^−1.7^, using the same staircase procedure (**Experiment 1b**) for 250 trials.

Naïve and inexperienced observers tested in **Experiment 1** first obtained experience with segmenting both kinds of boundaries over two training sessions prior to the experiment. During the first training session, they ran two full threshold series for segmenting both LTBs (*π_U_* cue) and LSBs (*C*_*M*_ cue). During the second training session, they ran one more series for both cues. Immediately after the second training session, they ran a final (4^th^) threshold series to estimate stimulus levels for each cue leading to JND (75% correct) performance.

#### Experiment 2: LTBs with constant luminance difference

In order to test the hypothesis that the key variable determining LTB segmentation performance is luminance difference, we generated a series of LTB stimuli having constant luminance difference arising from a fixed number (N = 8) of unbalanced (opposite color) micropatterns on opposite sides of the boundary. By adding an equal number of luminance-balanced micropatterns (i.e. having the same color) to both sides of the boundary (N = 0, 8, 16, 24, 32), we decreased the proportion of unbalanced micro-patterns, making the boundary more difficult to segment, while maintaining constant luminance difference across the boundary. Examples of such stimulus images with 0, 16 or 32 additional balanced pairs of micro-patterns are illustrated in **Fig. 5a**.

#### Experiment 3: Segmenting LTBs with varying RMS contrasts

In order to test further whether total luminance difference was a strong predictor of LTB segmentation performance, we repeated Experiment 1 for a single density (32 micro-patterns per side) while varying the maximum luminance *A* of each micro-pattern with respect to the screen mid-point luminance (0.5). This was accomplished by setting the maximum amplitude of each micro-pattern to three different levels with respect to the mid-point. W/B micro-pattern amplitudes were set at *A* = +/− 0.1, +/− 0.25, +/− 0.4 with respect to the luminance mid-point of 0.5 (*C*_*M*_ = 2*A* = 0.2, 0.5, 0.8). This had the effect of creating a large range of luminance differences across the LTB, for the same micro-pattern density. Examples of such stimuli are shown in **Fig. 6a**.

#### Experiment 4: Segmenting LTBs while ignoring masking LSBs

Of particular interest for the current study is investigating the relationship between the mechanisms used to segment LTBs and those used to segment LSBs. If the mechanisms are fully distinct, an observer should have little difficulty in segmenting a superimposition of an LTB and an LSB (either of the same or different orientations), when instructed to segment using only the LTB cue. Conversely, identical or highly overlapping mechanisms would lead to profound impairment of performance.

To investigate this question, we ran an experiment (**Experiment 4**) using author CJD, two naïve experienced observers (EMR, KNB), and N = 6 naïve inexperienced observers. Observers were instructed to segment an LTB target using proportion of unbalanced patterns *π_U_* as the segmentation cue, where *π_U_* was presented at JND (75% correct) as measured for that observer (determined from **Experiment 1a)**. For some trials, a masking LSB (also presented at that observer’s JND), which observers were instructed to ignore, was added to the LTB. There were three kinds of trials in this experiment: 200 *neutral* trials where the LTB was presented in isolation, 200 *congruent* trials with the LTB target and masking LSB having congruent boundary orientation (both cues left or right-oblique: see **Fig. 2c**), and 200 incongruent trials with the LTB target and masking LSB having incongruent orientations (one cue left-oblique, the other right-oblique: see **Fig. 2d**). For the (200) congruent stimuli, in half of trials (100) the two stimuli were phase-aligned (**Fig. 2c**, *left*), and for the other half (100) they had opposite phases (**Fig. 2c**, *right*).

#### Experiment 5: Effects of LSB masker on LTB segmentation thresholds

In order to explore the robustness of LTB segmentation to supra-threshold LSB maskers, two naïve, experienced observers (KNB, ERM) and author CJD segmented LTBs with a super-imposed LSB masker presented at various multiples of the LSB segmentation threshold (2x, 4x, 8x), yielding a masking LSB whose orientation was clearly visible. LTB segmentation thresholds were measured using the same staircase procedure as in **Experiment 1a**.

### Data Analysis

#### Psychometric function fitting

Data was fit using a signal-detection theory (SDT) psychometric model (Kingdom & Prins, 2016), where the proportion correct responses (*P*_*C*_) for a single-interval classification (1-AFC) task is given by

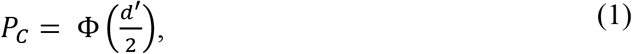

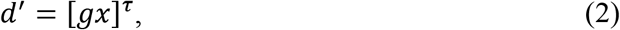

where *d*′ is the separation of the (unit variance) signal and noise distributions, with stimulus intensity *x*, and free parameters of gain *g* and transducer exponent *τ*. The SDT model was fit to psychophysical data using MATLAB® routines from the Palemedes Toolbox (http://www.palamedestoolbox.org/), as described in (Kingdom & Prins, 2016). Data was fit both with and without lapse rates, and nearly identical threshold estimates were observed in both cases, although sometimes fitting without lapse rates under-estimated the psychometric function slope. For the case of the model fitted using lapse rates,

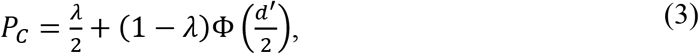

where λ denotes the lapse probability, which was constrained to lie in the range [0, 0.1].

#### Psychometric functions fit to luminance differences

Psychometric functions were also fit using one or two quantities computed from stimulus images. Given stimulus levels *x* used to generate the stimulus, we computed from each of the resulting images two quantities: *L*(*x*), which is the absolute value of the difference in luminance across the diagonal corresponding to the target orientation, and *C*(*x*), which is the global RMS stimulus contrast.

We then fit an alternative SDT psychometric function, where

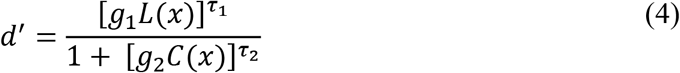

to model effects of global stimulus contrast *C*(*x*) that might co-vary with luminance differences *L*(*x*) as stimulus level *x* is varied. This model (4) is only appropriate for experiments in which the global stimulus contrast *C*(*x*) varies, since otherwise it is over-parametrized, and in these cases we set *g*_2_ = 0.

#### Image-computable model with one filtering stage

By design of the stimuli used in **Experiments 1-3**, for each trial image there is no difference in luminance across the anti-diagonal (the axis orthogonal to the stimulus orientation). Therefore, there was usually no need to take this into account when applying the model (4). However, in the masking experiment (**Experiment 4**), in the case where the masking LSB has an incongruent orientation, there will be a luminance difference across the anti-diagonal, which can potentially influence the decision. To analyze this data, we apply a slightly different model. In this model, illustrated schematically in **Fig. 4a**, we assume that each stimulus *x* gives rise to a decision variable *u*(*x*) which serves as input to the unit normal cumulative density function (CDF) Φ, so that the probability of a “right-oblique” behavioral response (*b* = *R*) is given by

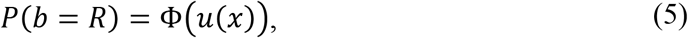

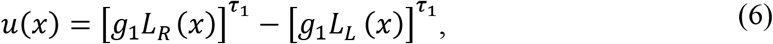

where *L*_*R*_(*x*), *L*_*L*_(*x*) are the absolute values of the luminance differences across the right- and left-diagonals. We also extended the model (6) to include divisive normalization by global stimulus contrast *C*(*x*), as in (4).

#### Image-computable model with two filtering stages

Masking data from **Experiment 4** was modeled using a two-stage model, illustrated in **Fig. 9a**. This model first convolves the image with on-center and off-center Difference-of-Gaussians (DOG) filters. The output of this first filtering stage is rectified and then passed to a second stage of filtering which computes a difference in first-stage activity across the left and right oblique diagonals. Second-stage filters were assumed to take a half-disc shape, integrating uniformly across the first stage outputs. The outputs of these second-stage filters are then subtracted to calculate a decision variable *u*(*x*). We fixed the first-stage DOG filter properties so that the standard deviation of the Gaussian defining the filter center is matched to the radius of the dots, while that defining the surround has a standard deviation twice that of the center. This choice is consistent with previous classification image studies of Gaussian detection in noise (Eckstein, Shimozaki & Abbey, 2002). Mathematically, this filter is defined as

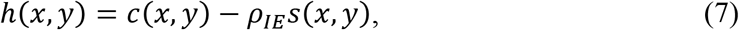

where *C*(*x*, *y*) denotes the center, and *s*(*x*, *y*) the surround, evaluated at (*x*, *y*). The only free variable for the first stage which we estimate from the data is the ratio *ρ_IE_* of the amplitudes of the center and surrounds, with *ρ_IE_* = 0 indicting no surround. If the rectified luminance differences (with nonlinear exponent *τ*_1_) from the left and right ON-center filters is given by 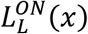, 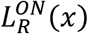, and from the OFF-center filters 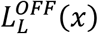, 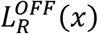, our decision variable is

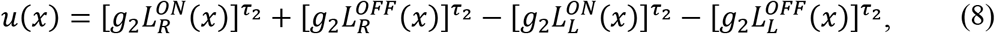

where *g*_2_, *τ*_2_ are gains and nonlinearities for the second-stage filters. The two-stage model only contains 4 free parameters (*ρ_IE_*, *τ*_1_, *τ*_2_, *g*_2_) which we estimate by fitting to data. To make computations tractable, we pre-filtered the stimuli with the center-surround DOG filters with IE amplitude ratios given by *ρ_IE_* = 0, 0.05, 0.1, 0.15, 0.2, 0.25, 0.3, 0.35, 0.4 and then optimized (MATLAB *fmincon*) the remaining parameters for each value of *ρ_IE_*. Initial starting points for the optimization were found using a 3-D grid search with *τ*_1_, *τ*_2_ taking grid values [0.5, 1, 2] and *g*_2_ taking grid values from 10^−3^ to 10^1^ in 5 log steps.

#### Bootstrapping psychometric functions

Bootstrapping was employed to determine the 95% confidence intervals for both the psychometric function thresholds (**Experiment 1**), as well as the proportion of correct responses predicted as a function of the stimulus level defined as either *π_U_* or absolute luminance difference (**Experiment 3**). For bootstrapping analyses, N = 100 or N = 200 simulated datasets were created as follows: For each stimulus level with *n*_*i*_ presentations and *c*_*i*_ experimentally observed correct responses (proportion of correct responses *p_i_* = *c_i_*/*n_i_*), we sampled from a binomial distribution having *ni* trials with probability *p*_*i*_ to create a simulated number of correct responses for that stimulus level. We fit our models to each of these simulated datasets, and obtained distributions of the psychometric function parameters, as well as the stimulus levels corresponding to JND (75% correct) performance, with confidence intervals being calculated using the standard deviation of the bootstrapped distributions.

## RESULTS

### Luminance texture boundary stimuli

In order to quantitatively examine the segmentation of luminance texture boundaries (LTBs), we defined a set of LTB stimuli which allowed us to vary the luminance across a boundary by varying the proportion of black and white micro-patterns within in each region (**Fig 2a**). When there are equal numbers of black (B) and white (W) micro-patterns on each side of the boundary, each micro-pattern is *balanced* by another of the same color on the other side. In this case, the luminance difference between regions is zero. When one side has more W patterns, and the opposite side has more B patterns, a proportion of the patterns on each side are *imbalanced*, giving rise to a difference in luminance across the diagonal. Therefore, we can modulate the luminance difference and therefore the boundary salience by changing the proportion of patterns on each side that are unbalanced (*π_U_*), as illustrated in **Fig 2a**. A value of *π_U_* = 0 corresponds to no boundary, whereas *π_U_* = 1 means that all the patterns on each side are the same.

Since both W and B micro-patterns have the same amplitude relative to the gray mid-point, the stimulus RMS contrast remains constant as we vary *π_U_*. Furthermore, when generating these stimuli we made sure that for each individual image there was no luminance difference across the orientation orthogonal to the boundary (the *anti-diagonal*). This ensured that there was no segmentation cue available which could mislead the observer to incorrectly classify the boundary as being in the opposite category.

### Experiment 1: Measuring segmentation thresholds

In **Experiment 1a**, we examined the ability of N = 17 observers (16 naïve, 14/16 inexperienced) to segment LTBs using the proportion of unbalanced micro-patterns (*π_U_*) as a cue. **Fig. 3a** shows the psychometric functions of two representative inexperienced observers (EMW, MCO) and two experienced observers (ERM, KNB). Nearly identical threshold estimates were obtained with and without lapse rates (**Supplementary Fig. S2a**). A histogram of JND thresholds (75% correct) for all observers is shown in **Fig. 3b.** The median observer could perform the task at with a threshold of *π_U_* = 0.31, and the best observer could reliably segment at *π_U_* = 0.16, suggesting a strong sensitivity to the proportion of unbalanced micro-patterns on the two surfaces.

**Figure 3:**
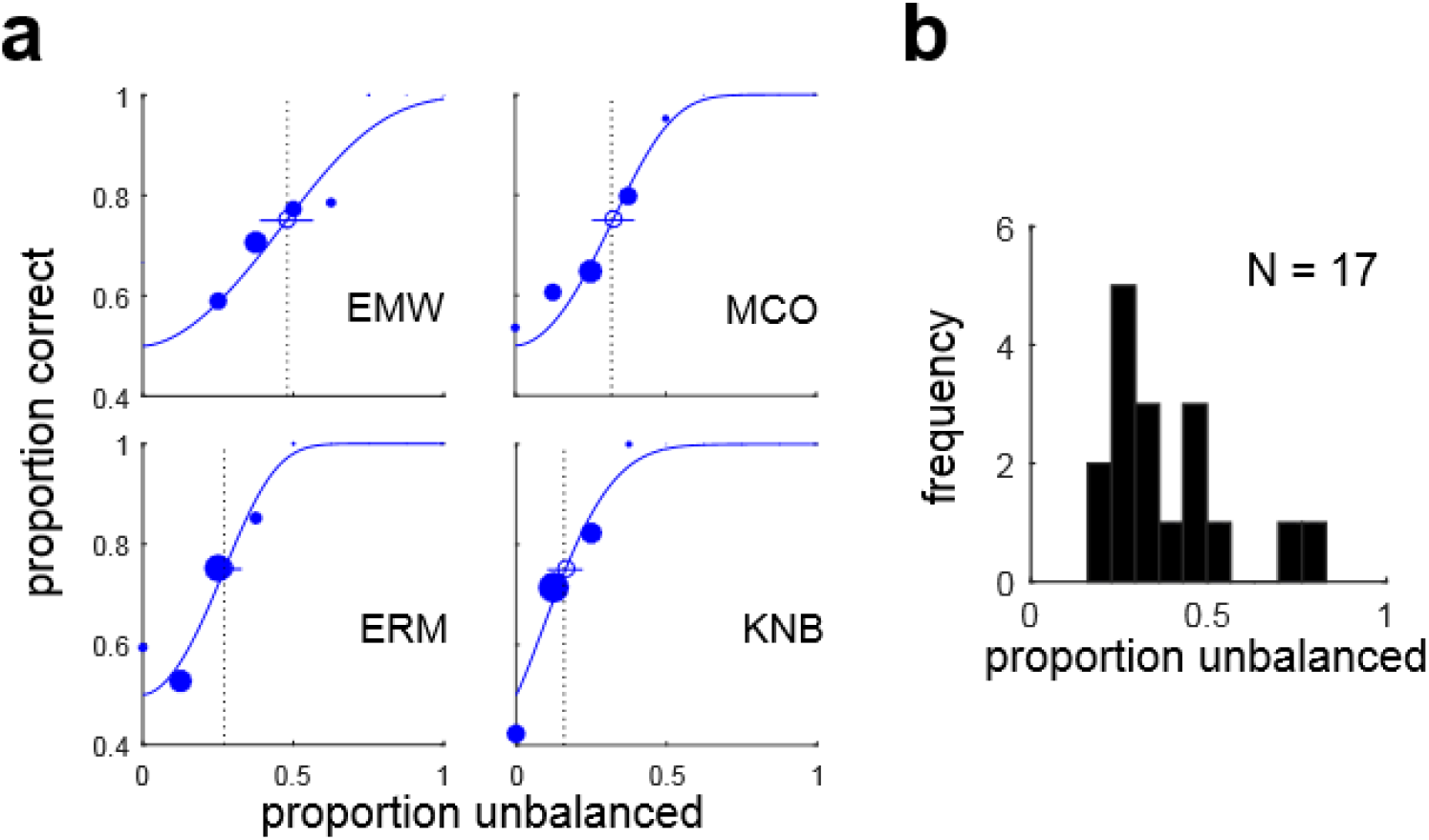
Psychometric functions and threshold distributions. (**a**) Psychometric functions and fitted functions based on SDT model (blue curves) for four observers (EMW, MCO, ERM, KNB) performing luminance texture boundary (LTB) segmentation (**Experiment 1a**) as a function of the proportion unbalanced micropatterns (*π_U_*), i.e. the proportion of micropatterns not having an opposite-polarity counterpart on the same side of the boundary. The size of each solid dot is proportional to the number of trials obtained at that level, and dashed black lines denote 75% thresholds for the fitted curves. Circles and lines indicate threshold estimates and 95% confidence intervals obtained from 200 bootstrapped re-samplings of the data. (**b**) Histogram of segmentation thresholds (*π_U_*) measured from all observers (N = 17) in **Experiment 1a**.

In **Experiment 1b** we also determined LSB segmentation thresholds for luminance disc stimuli like that shown in **Fig. 2b** in units of Michaelson contrast for the same N = 17 observers tested in **Experiment 1a** (**Supplementary Fig. S3**) Across the population of observers (**Supplementary Fig. S4**), we observed a significant positive rank-order correlation between LTB and LSB thresholds obtained in **Experiments 1a** and **1b** (Spearman’s ρ = 0.56; *p* = 0.019).

### Evaluating a simple model

One simple explanation for LTB segmentation performance is that the visual system is performing a simple luminance difference computation. As the proportion of unbalanced micro-patterns increases, so does this luminance difference, making the LTB more visible. We implemented an image-computable model like that shown in **Fig. 4a**, comprised of a single filtering-stage in which a left-oblique filter and right-oblique filter compute luminance differences across their respective boundaries, and the rectified, exponentiated outputs of these filters are subtracted to determine the probability the observer makes a “right-oblique” (R) response (**Eq. 4**). We see in **Fig. 4b** that this simple model predicts observer performance quite well as function of the luminance difference for LTB stimuli. Likewise, this model predicts performance well for LSB stimuli (**Supplementary Fig. S5**).

**Figure 4:**
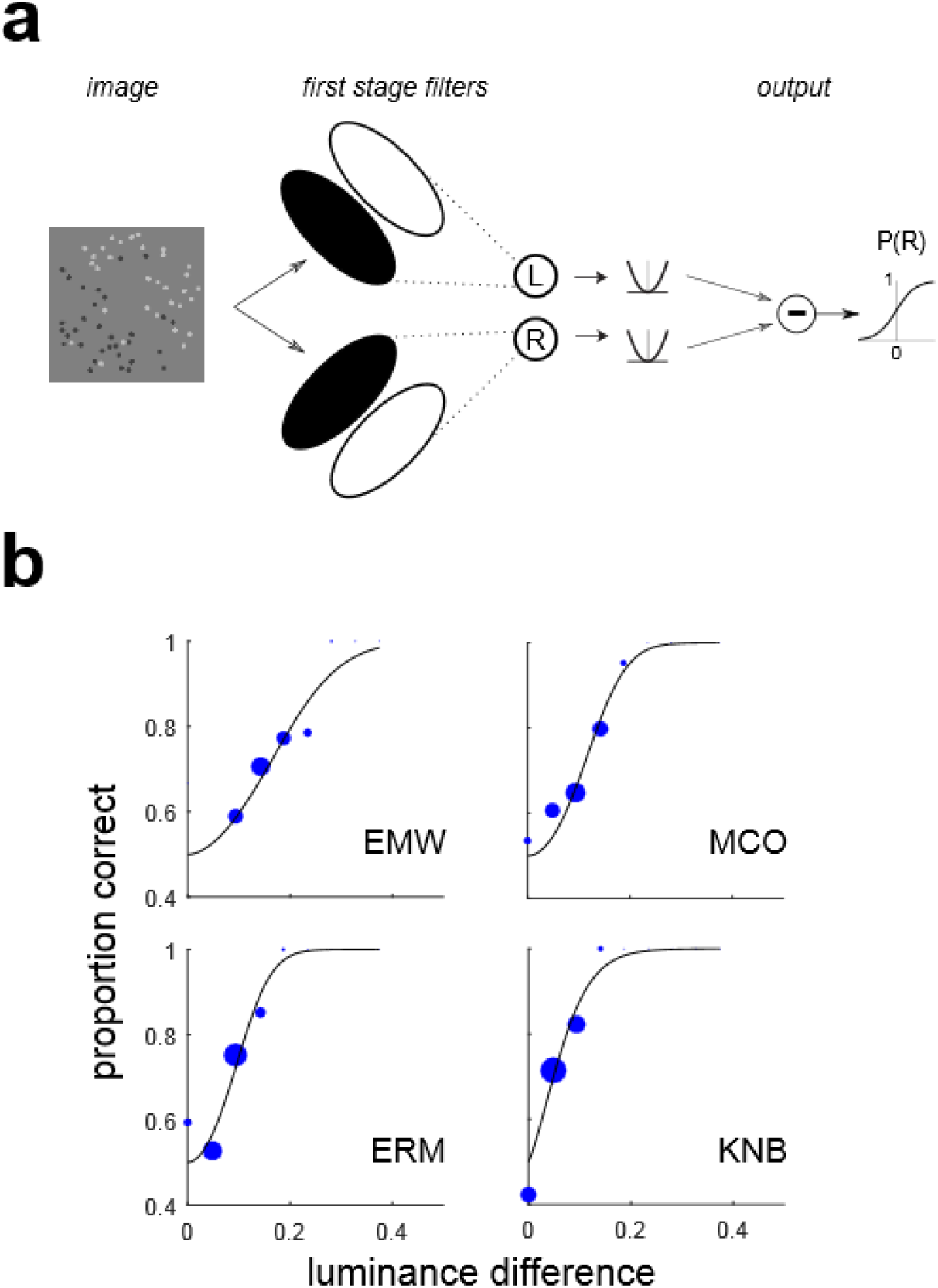
Single-stage filter model. (**a**) Model with a single stage of filtering. Luminance differences are computed across the left-oblique and right-oblique diagonals, passed through a rectifying, exponentiating nonlinearity and subtracted to determine the probability P(R) of observer classifying the boundary as right-oblique. (**b**) Fits of the model in (a) to LTB segmentation data from **Experiment 1a** for the same observers as in **Fig. 3a**.

### Experiment 2: Holding luminance difference constant

In order to directly test whether a simple luminance difference computation like that shown in **Fig. 4a** is adequate to explain LTB segmentation, in **Experiment 2,** we constructed a series of LTB stimuli having an identical number of unbalanced micro-patterns on each side, which provide the segmentation cue, while increasing the number of balanced patterns on each side, which serve as distractors. Stimuli from this experiment are illustrated in **Fig. 5a**. We see in **Fig. 5b** that for all three observers tested, performance decreases as the number of distractors increases, with all observers showing a significant effect of the number of distractors (Pearson’s chi-squared test; CJD: χ^2^(4) = 25.32, *p* < 0.001, ERM: χ^2^(4) = 34.817, *p* < 0.001, KNB: χ^2^(4) = 18.56, *p* = 0.001). These results argue against the hypothesis that LTB stimuli are segmented using a simple luminance difference computation, at least in cases like this where the total number of micro-patterns co-varies with the proportion of unbalanced patterns.

**Figure 5:**
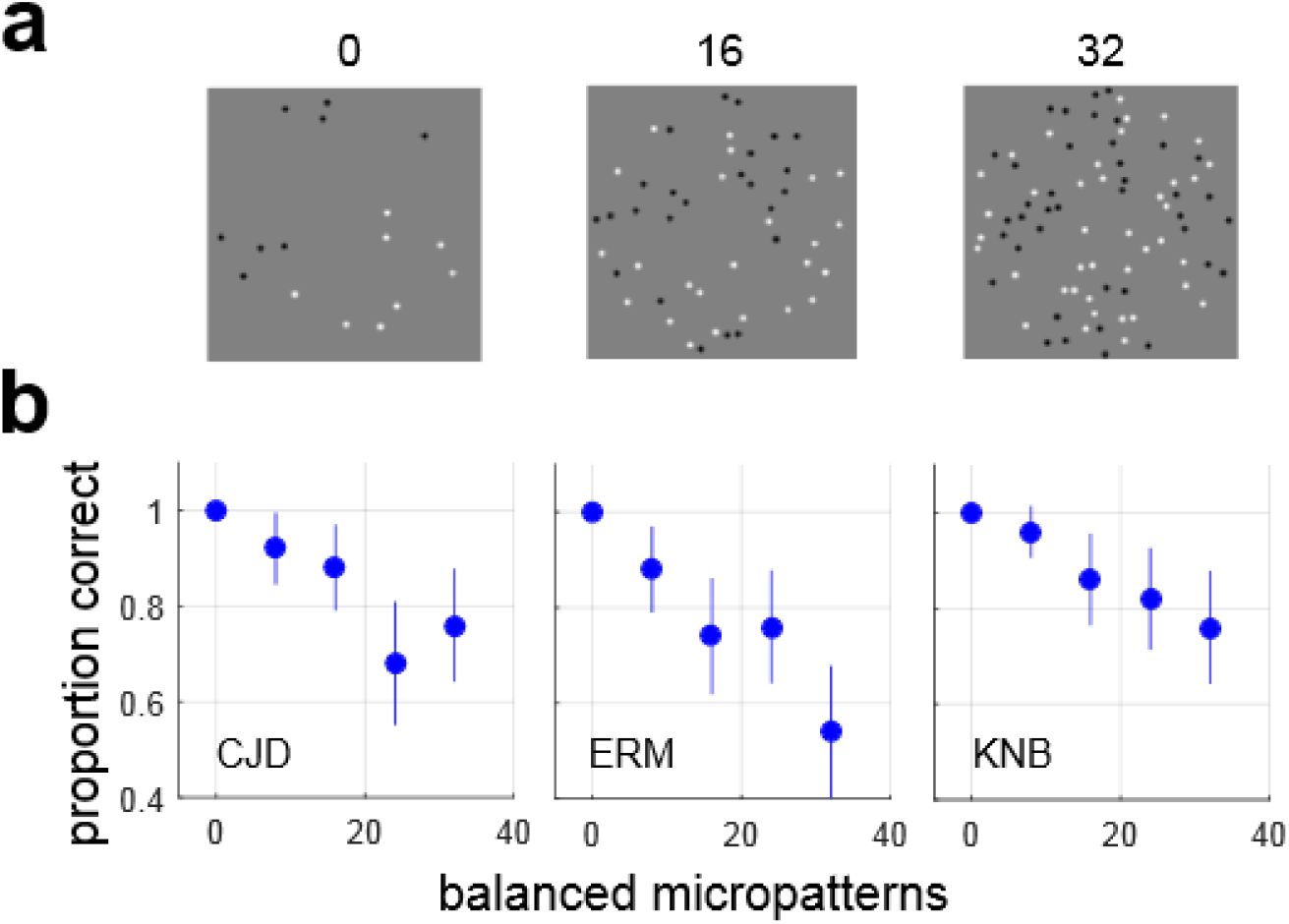
Holding luminance difference constant. (**a**) Examples of LTB stimuli used in **Experiment 2**, having an equal number (8) of unbalanced micropatterns on each side of the boundary, with varying numbers (0, 16, 32) of balanced micro-patterns. In this series, the luminance difference across the boundary is constant for all stimuli. (**b**) Proportion correct responses for three observers for differing numbers of balanced micropatterns. Lines indicate 95% binomial proportion confidence intervals for each level (N = 50 trials at each level). We see that performance degrades significantly with increasing numbers of balanced micropatterns, despite constant luminance difference. This suggests that a simple luminance difference computation may be inadequate to explain segmention of LTB stimuli.

### Experiment 3: Varying contrast while segmenting by proportion unbalanced patterns

As suggested by **Experiment 2**, a simple luminance difference computation is not a plausible candidate for segmenting LTB stimuli. In **Experiment 3**, we adduce additional evidence against this simplistic model. In this experiment, three observers (CJD, KNB, ERM) segmented LTB stimuli using the proportion of unbalanced micro-patterns *π_U_* as a cue, as in **Experiment 1a**. This was performed for three different levels of the stimulus Michaelson contrast (*C*_*M*_ = 0.2, 0.5, 0.8). This had the effect of creating drastically different regional luminance differences for stimuli in different series having the same proportion of unbalanced micro-patterns *π_U_* (**Fig. 6a**). As we see in **Fig. 6b**, *π_U_* (left panels) is a much better predictor of observer performance than the absolute luminance difference (right panels). Therefore, despite wide variation in the absolute difference in luminance across the boundary at different contrasts, observers are still able to detect differences in the proportion of light and dark areas in the two regions.

**Figure 6:**
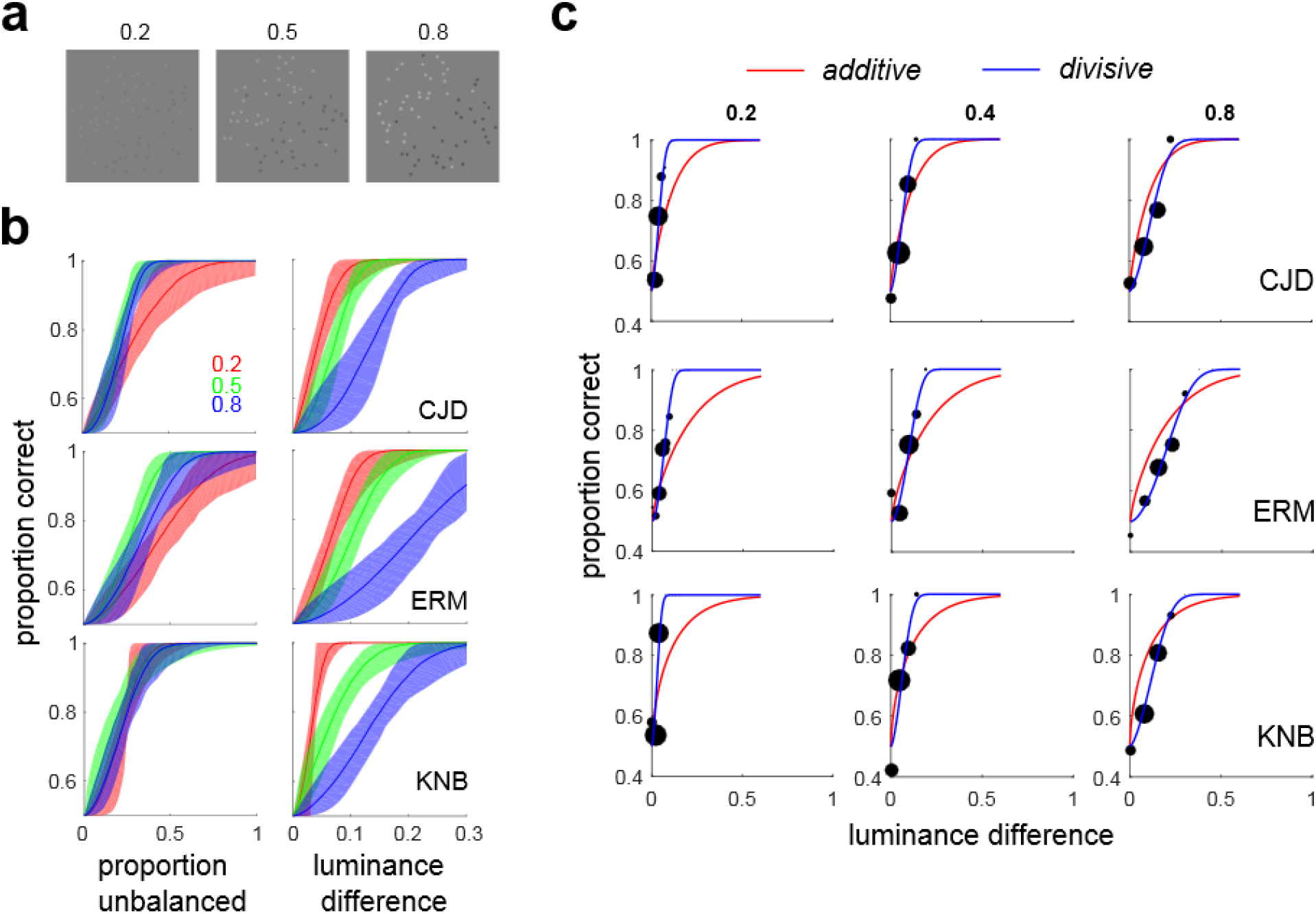
Using micro-pattern amplitude to vary global luminance difference. (**a**) Examples of LTB stimuli used in **Experiment 3**, with different Michaelson contrasts 0.2, 0.5, 0.8. (**b**) Bootstrapped SDT psychometric function fits (200 bootstrapped re-samplings) with 90 percent confidence intervals of observer performance as a function of proportion unbalanced micropatterns (left panels) and absolute luminance difference (right panels). This shows that identical luminance differences give rise to significantly different levels of observer performance for the three Michaelson contrasts (right panels), i.e. global luminance difference is a very poor predictor of performance. Instead, observer performance is much better predicted by the proportion of unbalanced micro-patterns, (almost) irrespective of micro-pattern amplitude (left panels). (**c**) Data from **Experiment 3** (black dots) and fits of the additive (red) and divisive (blue) signal detection theory models to the data. Each observer was tested at three different maximum micro-pattern amplitudes, which correspond to different Michaelson contrasts (0.2, 0.4, 0.8) of the stimuli. We see that a model incorporating a global luminance difference computation followed by contrast normalization (blue) provides an excellent fit to this data.

### Extending the one-stage model: Divisive computations

One can account for observer performance in **Experiments 2** and **3** using a single-stage model like that in **Fig. 4a** by introducing a contrast normalization operation (**Eq. 4**). Pooling data from all three contrast levels in **Experiment 3**, we fit both the standard SDT model (**Eq. 2**) using simple luminance difference only, as well as the divisive SDT model (**Eq. 4**) incorporating both luminance difference and RMS contrast normalization. As we see in **Fig. 6c**, the fit of the standard additive SDT model (red lines) is quite poor compared to the divisive SDT model (blue lines). Since the divisive model has more parameters, we compare the goodness-of-fit using the Bayes Information Criterion (BIC), which rewards goodness of fit while penalizing model complexity (Schwarz, 1978; Bishop, 2006). The BIC analysis suggests a strong preference (Kaas & Raferty, 1995) for the divisive model for all observers (**Supplementary Table S1**). Similar results were obtained using models with lapse rates estimated as well (**Supplementary Fig. S6**). In addition, we see that the divisive SDT model (**Eq. 4**) is able to do a reasonably good job of predicting observer performance in **Experiment 2** (**Supplementary Fig. S7,***red symbols*).

### Experiment 4: Segmenting LTBs while ignoring LSBs

The results of Experiments 1-3 suggest that a model implementing a luminance difference computation (**Fig. 4a**) with contrast normalization can potentially explain LTB segmentation performance. However, one weakness of a single-stage model computing simple luminance differences is that it may be susceptible to interference from masking LSBs having incongruent orientations. Motivated by these considerations, in **Experiment 4** we investigated the extent to which segmentation of LTB stimuli is influenced by the presence of masking LSB stimuli which observers are instructed to ignore. The logic of this paradigm is that if LTBs and LSBs are processed by entirely different mechanisms, then the presence of a task-irrelevant LSB should have no effect on segmentation using the LTB cue. If one cue cannot be ignored, it suggests that there may be some overlap or interaction between the mechanisms. This sort of paradigm was used in a previous study (Saarela & Landy, 2012) to demonstrate that second-order color and texture cues were not processed independently.

In **Experiment 4**, N = 9 observers segmented LTB stimuli as in **Experiment 1a** using proportion of unbalanced micro-patterns as a cue, with *π_U_* set to the observer’s 75% performance threshold. For 200 *neutral* trials, the LTB was presented in isolation, for 200 *congruent* trials a masking LSB at segmentation threshold was presented with the same orientation (L/R oblique) as the target (**Fig. 2c**), and for 200 *incongruent* trials the LSB was presented at the orthogonal orientation (**Fig. 2d**). For half of the congruent trials, the LTB and LSB were phase-aligned (**Fig. 2c, “con-0”, left**), and for the other half they were opposite-phase (**Fig. 2c, “con-180”, right**).

As we can see from **Fig. 7a**, performance when segmenting LTB stimuli when using *π_U_* as the cue is quite robust to interference from masking LSB stimuli. Statistical tests (Pearson’s Chi-squared) comparing observer performance across all three conditions did not find any significant effect of condition (*neutral* (**neu**), *congruent* (**con**), *incongruent* (**inc**)) for any individual observer (**Supplementary Table S2**). Pooling across all observers, we did however obtain significantly different (**χ^2^**(2) = 15.319, *p* < 0.001) values of proportion correct for each condition (**neu**: 0.8217, **con**: 0.8622, **inc**: 0.8189), due to slightly enhanced performance for congruent masking LSBs, since there was no impairment for incongruent masking LSBs (**χ^2^**(1) = 0.047, *p* = 0.828). The enhanced performance for congruent masking LSBs was phase-dependent, as seen in **Fig. 7b**. For the aligned-phase case (**con-0**), we observe significant improvements in performance over the neutral condition for 4/9 observers (**Supplementary Table S2**). We fail to find any significant difference in individual observer’s performance between the neutral and opposite-phase (**con-180**) cases. Pooling across observers, we find significant differences (**χ^2^**(1) = 24.383, *p* < 0.001) between the proportions correct for the neutral case and the aligned-phase case (**neu:**0.8217, **con-0:**0.8944). However, we fail to find a significant difference (**χ^2^**(1) = 0.288, *p* = 0.592) between the proportion correct in the neutral case and the opposite-phase case (**con-180:**0.8300). In at least some observers (3/9 total, 2/8 naive) we see improved performance for phase-aligned compared to opposite-phase boundaries in the congruent case (**Fig. 7b**, **Supplementary Table S3)**, as well as a significant effect (**χ^2^**(1) = 15.732, *p* < 0.001) pooling across all observers (**con-0**: 0.8944, **con-180**: 0.8300).

**Figure 7:**
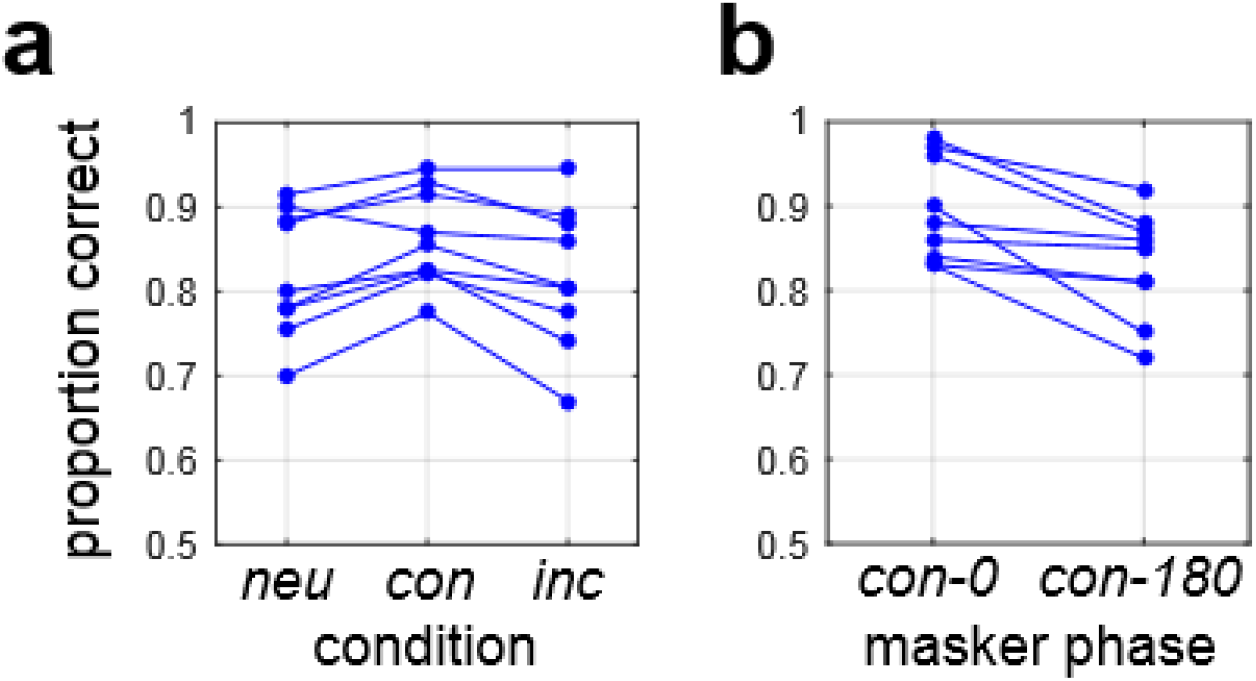
Effects of masking LSBs on LTB segmentation. (**a**) Performance for N = 9 observers in **Experiment 4**, segmenting LTB stimuli using a proportion of unbalanced micro-patterns (*π_U_*), set at 75% JND for each observer, as measured in **Experiment 1a**. We see similar performance for most observers in the absence of a masker (neutral case, **neu**) as well as with a masker having congruent (**con**) and incongruent (**inc**) orientation. Here the congruent case pools across in-phase and opposite-phase conditions. (**b**) Performance for same observers for congruent stimuli which are in-phase (**con-0**) and opposite-phase (**con-180**).

In **Experiment 4**, observers segmenting LTBs using proportion unbalanced patterns as a cue were relatively unimpaired by the presence of masking LSBs having an incongruent orientation, at least when the LSBs were presented at their segmentation thresholds. In **Experiment 5**, we studied the effects of supra-threshold LSB maskers on LTB segmentation in three experienced observers (ERM, KNB, CJD). Consistent with **Experiment 4**, we found that although LTB maskers presented well above threshold can somewhat raise LSB segmentation thresholds, this effect was generally modest (**Supplementary Fig. S8**).

### Evaluating one-stage and two-stage models

Given our findings that LTB segmentation is fairly robust to interference from masking LSB stimuli, it seemed likely that LTBs might be detected by a distinct mechanism. Consequently, we considered the possibility that LTB segmentation may be better explained by a model like that shown in **Fig. 8a** with two stages of processing, rather than a single stage as in the model in **Fig. 4a**. The first stage is comprised of small-scale spatial filters, implemented here as center-surround filters (see Methods), which are convolved with the input image and whose outputs are passed through a rectifying nonlinearity. The second stage analyzes the first-stage outputs, with two large-scale filters selective for left-oblique and right-oblique boundaries. These second-stage filter outputs are rectified and subtracted to determine the probability of an “R” response. Note that since the center-surround filters in the first stage are poorly driven by constant light levels, this model can in principle exhibit robustness to interference from LSBs, while still permitting some degree of influence, depending on the relative strengths of the center-surround units, which determines the response of the filter to mean luminance.

**Figure 8:**
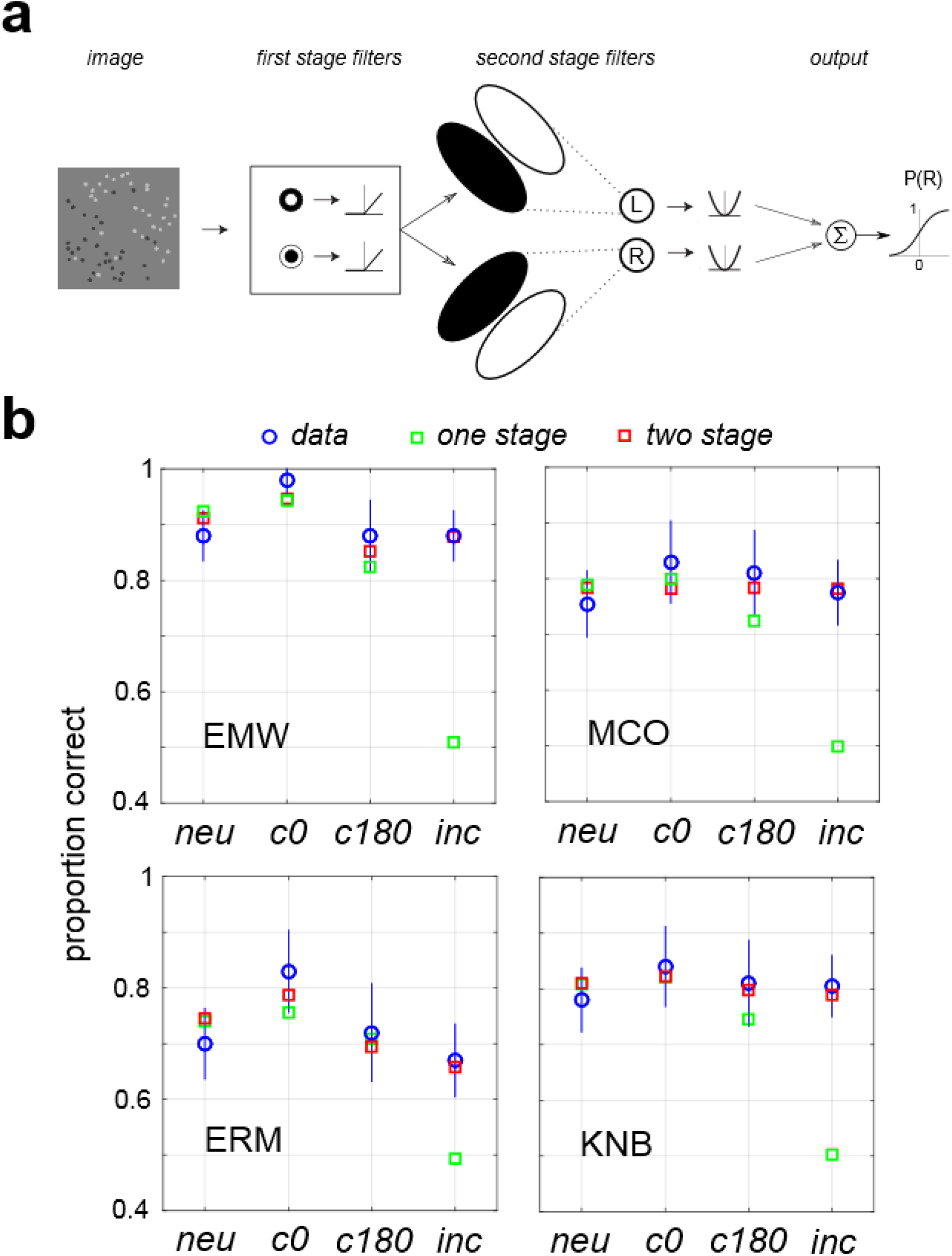
Two-stage model fits Experiment 4 results. (**a**) Model with two cascaded stages of filtering. The first stage of this model detects texture elements (here, micro-patterns) on a fine spatial scale. The second stage looks for differences in the outputs of these first-stage filters on the coarse spatial scale of the texture boundary, at either of two possible orientations. Such a model can detect differences in the proportions of black and white micro-patterns on opposite sides of the boundary, while being fairly robust to interference from luminance steps. (**b**) Fits of single-stage model (green squares) and two-stage model (red squares) to data from **Experiment 4** (blue circles, lines denote 95% confidence intervals), for four ways of combining LTB and LSB stimuli: neutral (**neu**); congruent, in-phase (**c0**); congruent, opposite phase (**c180**); and incongruent (**inc**).

**Fig. 8b**. shows the fits of both the one-stage model (**Fig. 4a**) and two stage model (**Fig. 8a**) to data obtained from **Experiment 4** for four observers (EMW, MCO, ERM, KNB). One stage models were fit both with and without divisive normalization terms, and identical predictions of observer performance were obtained. We see in **Fig. 8b** that although both one-stage (green squares) and two-stage (red squares) models fit observer performance (blue circles) in the neutral (**neu**) and two congruent cases, the one-stage model clearly fails to account for observer performance in the incongruent case (**inc**), predicting near-chance performance. Plots like those in **Fig. 8b** are shown for all other observers in **Supplementary Fig. S9**. The lack of robustness of the one-stage model to incongruently oriented LSBs argues strongly in favor of the two-stage model as a more plausible mechanism for LTB segmentation, at least in the presence of interfering LSBs. Fitting these two models to all observers in **Experiment 4** and plotting the preference for the two-stage model (BIC_2_ – BIC_1_, **Supplementary Fig. S10a**) reveals over the set of N = 9 observers a significant preference for the two stage model (single sample t-test, mean = 30.22, *t*(8) = 4.077, *p* < 0.004).

As shown in **Fig. 8b**, for the majority of observers, we obtain better LTB segmentation performance in the presence of a congruent boundary with aligned phase (**con-0**) than opposite phase (**con-180**). This difference is also evident for some of the other observers (**Supplementary Fig. S9**) Interestingly, the two stage model allows for LSB stimuli to potentially influence LTB segmentation via a center-surround imbalance of the first-stage filters which can provide a mean-luminance ("DC") response. That is, if the on-center (off-center) filters have a small positive (negative) response to constant light levels, this would allow LSB stimuli to exert an excitatory influence on the second-stage filters, potentially explaining the slightly improved performance for the phase-aligned versus opposite-phase congruent case in **Experiment 4** (**Fig. 7b, 8b**). Over the population of observers (**Supplementary Fig. S10b**), we found the fitted first stage on-center filters all had a positive DC response (single-sample t-test, mean = 6.91, *t*(8) = 2.92, *p* < 0.019). Finally, we investigate whether the two-stage model in **Fig. 8a** can also account for the results of **Experiment 3** (**Fig. 6**). We find that as with the one-stage model, an excellent fit to the data (blue lines) is obtained using the two-stage model when a divisive normalization term is included (**Supplementary Fig. S11**).

## DISCUSSION

### Summary

Over half a century of research in modern vision science and has investigated visual texture segmentation using parametric stimuli (Julesz, 1962, 1981; Landy, 2013; Victor, 2017). However, this psychophysical work has largely focused on manipulating second-order and higher-order statistical properties which characterize textures, while holding first-order (luminance) cues constant (e.g., Zavitz & Baker, 2013, 2014). This is a sensible research strategy because it neatly isolates the problem of understanding how higher-order statistics influence segmentation. However, it is ultimately incomplete since natural region boundaries typically contain first-order cues like color and luminance (Johnson & Baker, 2005; Ing, Wilson, & Geisler, 2010; Mely et al., 2016; Breuil et al., 2019), which are known to combine with higher-order cues for localization and segmentation (Rivest & Cavanaugh, 1996; McGraw, Whitaker, Badcock, & Skillen, 2003; Ing et al., 2010; DiMattina et al., 2012). In most studies in which first-order cues are manipulated they are presented as steps or gratings (e.g., Elder & Sachs, 2004; McIlhagga, 2018), or when they are measured from natural images, it is as average luminance within a region (Ing et al., 2010; DiMattina et al., 2012). However, as we see in **Fig. 1**, differences in mean luminance can also be caused by differences in the proportion of light and dark pixels in each surface region, with no abrupt change in albedo at the boundary. We refer to boundaries of this kind as luminance texture boundaries (LTBs), to distinguish them from luminance step boundaries (LSBs). Understanding whether or not these two kinds of luminance cue (LTB, LSB) are processed via the same, different, or partially overlapping mechanisms is of great utility for understanding how first-order and higher-order cues combine to enable natural boundary segmentation. The present study provides a first step in this direction, suggesting that multiple mechanisms may contribute to luminance-based boundary segmentation in natural vision.

### Multiple mechanisms for segmentation using luminance cues

Clearly, whenever there are mean differences in luminance between two regions, a single stage of linear filtering (**Fig. 4a**) is capable of detecting this difference, for both LTBs (**Fig. 4b**) and LSBs alike. However, this simplistic model would make the prediction that for any two boundaries with equal luminance differences, segmentation performance should be identical. Explicitly testing this idea in **Experiment 2** and **Experiment 3** lead us to reject this model. Further exploration revealed that we can however explain the LTB segmentation data in **Experiments 2, 3** with a single stage of linear filtering if we incorporate a divisive operation (Carandini & Heeger, 2012) which normalizes filter outputs by global RMS contrast. Nevertheless, even with this improvement, any model positing a single stage of filtering computing a luminance difference is highly susceptible to interference from stimuli which provide extraneous luminance cues, for instance a shadow edge (LSB) with an orientation conflicting with the LTB orientation. We test this prediction explicitly in **Experiment 4**, where we investigated the ability of observers to segment LTB stimuli in the presence of masking LSB stimuli. In this experiment, we find that LTB segmentation is remarkably robust to interference from masking LSB stimuli. This robustness to masking argues against the idea that a single stage of filtering is adequate to fully explain LTB segmentation. Further investigation with supra-threshold LSB maskers (**Experiment 5**) added further support to the notion of separate mechanisms, although we did observe some degree of influence of LSB masking stimuli on LTB segmentation performance (**Supplementary Fig. S8**), as was also the case in **Experiment 4**.

We posit that two sequential stages of filtering on different spatial scales may be required to explain LTB segmentation, and implement the two-stage model is shown in **Fig. 8a**. It is comprised of an initial layer of filtering on a local spatial scale which detects the micro-patterns, followed by a second-stage of filtering which looks for spatial differences in the rectified outputs of the first-stage filters on a global scale. This model successfully explains the ability of observers to segment LTB stimuli in the presence of masking LSBs (**Fig. 8b**), and fits LTB segmentation data obtained in **Experiment 3** (**Supplementary Fig. S11**). Although the first stage filters in our model are implemented as center-surround filters, which are known to be present in area V1 (Ringach, Shapley, & Hawken, 2002; Talebi & Baker, 2012), orientation tuned mechanisms pooled across different orientations can in principle serve the same function (Motoyoshi & Kingdom, 2007). This general model architecture is known as the Filter-Rectify-Filter model (Chubb & Landy, 1991), and has been applied in dozens of studies to model texture segmentation and second-order vision (Landy, 2013). To our knowledge, the present study is the first time that it has been explicitly demonstrated that an FRF-style model can describe how observers segment textures defined entirely by first-order luminance cues.

One important finding from our psychophysical work is that although LTB segmentation is highly robust to interference from masking LSB stimuli, it is not entirely independent. For instance, in **Experiment 4** we found that segmentation performance was slightly better when the LTB an LSB having congruent orientation were phase-aligned compared to opposite phase (**Fig. 7b**). Furthermore, **Experiment 5** revealed higher LTB segmentation thresholds for supra-threshold LSB maskers, although this effect was very modest for two of the three observers tested. This interaction between LTB and LSB cues could arise in one of two possible ways. One possibility, suggested by our model fitting, is that the first-stage filters have a non-zero DC response. In particular, we observed that the on-center filters which best fit the data from **Experiment 4** had a slightly positive response to a constant uniform stimulus (**Supplementary Fig. S10**). This nonzero DC response is consistent with previous psychophysical studies (Eckstein et al., 2002), as well as known neurophysiology of center-surround retinal ganglion cells (Croner & Kaplan, 1995). However, another possibility is that the final decision arises by integrating the outputs of a two-stage model like that in **Fig. 8a** with zero DC response with the outputs of a single-stage model like that in **Fig. 4a**. Such a model would also be consistent with our observations, and it is of interest for future work to design an experiment which could distinguish between these two possibilities.

### Future directions

Although natural surfaces may have luminance differences which arise due to luminance texture boundaries, many other textural differences do not involve changes in luminance. Micro-pattern orientation, density, and contrast and others all provide powerful segmentation cues (Dakin & Mareschal, 2000; DiMattina & Baker, 2019; Zavitz & Baker, 2013, 2014; Wolfson & Landy, 1995; Motoyoshi & Kingdom, 2007), which must be combined with luminance cues to enable segmentation in natural vision. It is of great interest for future research to understand how luminance textures combine with other cues. In particular, one could define black and white micro-patterns as oriented bars instead of the dots used here, and simultaneously vary orientation and luminance cues to see how these cues summate, i.e. via probability summation or additive summation (Kingdom et al., 2015). Such an experiment would greatly expand the literature on the interaction of first-order and second-order cues, which has largely been limited to simple detection experiments in which the first-order cues were presented as gratings (Schofield & Georgeson, 1999; Allard & Faubert, 2007). Another interesting direction of research would be to consider how luminance steps and luminance textures combine for boundary segmentation. In many cases, two surfaces may have both kinds of cues defining luminance difference, and therefore combining both cues will be helpful for segmentation. Although we suggest that the mechanisms are not identical, they are most likely overlapping and therefore this kind of psychophysical summation experiment would be very interesting.

The present study strongly suggests the possibility of neural mechanisms tuned to LTBs which are minimally influenced by overlapping LSBs. We hypothesize that individual neurons tuned to LTBs will most likely be found in extra-striate areas, for instance V2 and V4, which are known to contain units sensitive to second-order boundaries (Mareschal & Baker, 1998; Schmid, Purpura, & Victor, 2014) and units exhibiting texture selectivity (Okazawa, Tajima, & Komatsu, 2017). As suggested by our psychophysical models, neurons at higher areas of the visual pathway may receive inputs from neurons in V1 or V2 responsive to the micro-patterns or texture elements. If the afferent presynaptic V1 neurons in one spatial region are optimally driven by light micro-patterns, and those in an adjacent spatial region prefer dark micro-patterns, the downstream extra-striate neuron will be sensitive to differences in the proportion in light and dark micro-patterns in these adjacent regions. It is of great interest for future neurophysiology studies to see if neurons can be observed which are selectively responsive to LTB stimuli, while being poorly driven, if at all, by step edges. Such neurons could provide a physiological basis for the ability to segment surface boundaries in the presence of shadows and distinguish shadow edges from boundaries (Vilankar, Golden, Chandler, & Field, 2014; Breuil et al., 2019).

Finally, a large body of work has demonstrated that deep neural networks trained on visual tasks like object recognition develop intermediate-layer representations which are sensitive to textural features (Kriegeskorte, 2015; Guclu & van Gerven, 2015). An entire sub-field of computational neuroscience known as “artificial neurophysiology” has developed to analyze the selectivity properties of units in these deep networks, and to interpret their results in light of known neurophysiology. It would be of great interest for future investigation to do an artificial neurophysiology study on deep neural networks resembling the ventral visual stream (Guclu & van Gerven, 2015) in order to look for neurons which are tuned to luminance texture boundaries while being relatively unresponsive to luminance steps, and to see if decoding such a population of units can account for human performance psychophysical in texture segmentation tasks.

## ACKNOWLEDGEMENTS

C.D. would like to thank the FGCU students in the Computational Perception lab for help with data collection.

## AUTHOR CONTRIBUTIONS

C.D. and C.B. conceptualized the study. C.D. created the stimuli, performed the experiments, and analyzed the data. C.D. and C.B. wrote and edited the manuscript.

## COMPETING INTEREST STATEMENT

The authors declare no competing interests.

## DATA AVILABAILITY

All data is available from author C.D. upon request.

## Notes

### Competing Interest Statement

The authors have declared no competing interest.

## REFERENCES

Allard, R., & Faubert, J. (2007). Double dissociation between first-and second-order processing. Vision Research, 47(9), 1129–1141.

Bishop, C. M. (2006). Pattern recognition and machine learning. Springer.

Brodatz, P. (1966). Textures: A photographic album for artists and designers. Dover Publications.

Brainard, D. H. (1997). The psychophysics toolbox. Spatial Vision, 10(4), 433–436.

Breuil, C., Jennings, B. J., Barthelmé, S., & Guyader, N. (2019). Color improves edge classification in human vision. PLoS Computational Biology, 15(10), e1007398.

Carandini, M., & Heeger, D. J. (2012). Normalization as a canonical neural computation. Nature Reviews Neuroscience, 13(1), 51–62.

Chubb, C., & Landy, M. S. (1991). Orthogonal distribution analysis: A new approach to the study of texture perception. In: Computational Models of Visual Processing (Eds: Landy, M.S. & Movshon, J.A.). MIT Press.

Croner, L. J., & Kaplan, E. (1995). Receptive fields of P and M ganglion cells across the primate retina. Vision research, 35(1), 7–24.

Dakin, S. C., & Mareschal, I. (2000). Sensitivity to contrast modulation depends on carrier spatial frequency and orientation. Vision Research, 40(3), 311–329.

DiMattina, C., & Baker Jr, C. L. (2019). Modeling second-order boundary perception: A machine learning approach. PLoS Computational Biology, 15(3), e1006829.

DiMattina, C., Fox, S. A., & Lewicki, M. S. (2012). Detecting natural occlusion boundaries using local cues. Journal of vision, 12(13), 15–15.

Eckstein, M. P., Shimozaki, S. S., & Abbey, C. K. (2002). The footprints of visual attention in the Posner cueing paradigm revealed by classification images. Journal of Vision, 2(1), 3–3.

Elder, J. H., & Sachs, A. J. (2004). Psychophysical receptive fields of edge detection mechanisms. Vision Research, 44(8), 795–813.

Güçlü, U., & van Gerven, M. A. (2015). Deep neural networks reveal a gradient in the complexity of neural representations across the ventral stream. Journal of Neuroscience, 35(27), 10005–10014.

Hansen, B. C., & Hess, R. F. (2006). The role of spatial phase in texture segmentation and contour integration. Journal of Vision, 6(5), 5–5.

Hubel, D. H., & Wiesel, T. N. (1962). Receptive fields, binocular interaction and functional architecture in the cat’s visual cortex. The Journal of Physiology, 160(1), 106.

Ing, A. D., Wilson, J. A., & Geisler, W. S. (2010). Region grouping in natural foliage scenes: Image statistics and human performance. Journal of vision, 10(4), 10–10.

Johnson, A. P., & Baker, C. L. (2005). Spatiochromatic statistics of natural scenes: first-and second-order information and their correlational structure. JOSA A, 22(10), 2050–2059.

Julesz, B. (1962). Visual pattern discrimination. IRE transactions on Information Theory, 8(2), 84–92.

Julesz, B. (1981). Textons, the elements of texture perception, and their interactions. Nature, 290(5802), 91–97.

Kass, R. E., & Raftery, A. E. (1995). Bayes factors. Journal of the american statistical association, 90(430), 773–795.

Kingdom, F.A.A., Baldwin, A. S., & Schmidtmann, G. (2015). Modeling probability and additive summation for detection across multiple mechanisms under the assumptions of signal detection theory. Journal of Vision, 15(5), 1–1.

Kingdom, F.A.A., & Prins, N. (2016). Psychophysics: a practical introduction. Academic Press.

Kriegeskorte, N. (2015). Deep neural networks: a new framework for modeling biological vision and brain information processing. Annual review of vision science, 1, 417–446.

Landy, M. S. (2013). Texture analysis and perception. In: The New Visual Neurosciences (Eds: Werner, J. S., & Chalupa, L. M.). MIT Press.

Leek, M. R. (2001). Adaptive procedures in psychophysical research. Perception & Psychophysics, 63(8), 1279–1292.

Mareschal, I., & Baker, C. L. (1998). A cortical locus for the processing of contrast-defined contours. Nature Neuroscience, 1(2), 150–154.

Marr, D. (1982). Vision: A computational investigation into the human representation and processing of visual information. Henry Holt and Co. Inc., New York, NY.

Martin, D. R., Fowlkes, C. C., & Malik, J. (2004). Learning to detect natural image boundaries using local brightness, color, and texture cues. IEEE Transactions on Pattern Analysis and Machine Intelligence, 26(5), 530–549.

McGraw, P. V., Whitaker, D., Badcock, D. R., & Skillen, J. (2003). Neither here nor there: Localizing conflicting visual attributes. Journal of Vision, 3(4), 2–2.

McIlhagga, W. (2018). Estimates of edge detection filters in human vision. Vision Research, 153, 30–36.

McIlhagga, W. H., & May, K. A. (2012). Optimal edge filters explain human blur detection. Journal of Vision, 12(10), 9–9.

McIlhagga, W., & Mullen, K. T. (2018). Evidence for chromatic edge detectors in human vision using classification images. Journal of Vision, 18(9), 8–8.

Mély, D. A., Kim, J., McGill, M., Guo, Y., & Serre, T. (2016). A systematic comparison between visual cues for boundary detection. Vision research, 120, 93–107.

Motoyoshi, I., & Kingdom, F. A. (2007). Differential roles of contrast polarity reveal two streams of second-order visual processing. Vision Research, 47(15), 2047–2054.

Okazawa, G., Tajima, S., & Komatsu, H. (2017). Gradual development of visual texture-selective properties between macaque areas V2 and V4. Cerebral Cortex, 27(10), 4867–4880.

Parker, A. J., & Hawken, M. J. (1988). Two-dimensional spatial structure of receptive fields in monkey striate cortex. JOSA A, 5(4), 598–605.

Pelli, D. G. (1997). The VideoToolbox software for visual psychophysics: Transforming numbers into movies. Spatial vision, 10(4), 437–442.

Ringach, D. L., Shapley, R. M., & Hawken, M. J. (2002). Orientation selectivity in macaque V1: diversity and laminar dependence. Journal of Neuroscience, 22(13), 5639–5651.

Rivest, J., & Cabanagh, P. (1996). Localizing contours defined by more than one attribute. Vision Research, 36(1), 53–66.

Saarela, T. P., & Landy, M. S. (2012). Combination of texture and color cues in visual segmentation. Vision Research, 58, 59–67.

Schmid, A. M., Purpura, K. P., & Victor, J. D. (2014). Responses to orientation discontinuities in V1 and V2: physiological dissociations and functional implications. Journal of Neuroscience, 34(10), 3559–3578.

Schofield, A. J., & Georgeson, M. A. (1999). Sensitivity to modulations of luminance and contrast in visual white noise: Separate mechanisms with similar behaviour. Vision Research, 39(16), 2697–2716.

Schwarz, G. (1978). Estimating the dimension of a model. The annals of statistics, 6(2), 461–464.

Talebi, V., & Baker, C. L. (2012). Natural versus synthetic stimuli for estimating receptive field models: a comparison of predictive robustness. Journal of Neuroscience, 32(5), 1560–1576.

Victor, J. D., Conte, M. M., & Chubb, C. F. (2017). Textures as probes of visual processing. Annual Review of Vision Science, 3, 275–296.

Vilankar, K. P., Golden, J. R., Chandler, D. M., & Field, D. J. (2014). Local edge statistics provide information regarding occlusion and nonocclusion edges in natural scenes. Journal of Vision, 14(9), 13–13.

Wolfson, S. S., & Landy, M. S. (1995). Discrimination of orientation-defined texture edges. Vision Research, 35(20), 2863–2877.

Zavitz, E., & Baker, C. L. (2013). Texture sparseness, but not local phase structure, impairs second-order segmentation. Vision Research, 91, 45–55.

Zavitz, E., & Baker, C. L. (2014). Higher order image structure enables boundary segmentation in the absence of luminance or contrast cues. Journal of Vision, 14(4), 14–14.

