## Supplementary-Material for "Segmenting surface boundaries using luminance cues: Underlying mechanisms"

**TABLE S1**

| | **BIC: Additive** | | **BIC: Divisive** | | **$BIC_D$ - $BIC_A$** | |
|---|---|---|---|---|---|---|
| | ***None*** | ***Lapse*** | ***None*** | ***Lapse*** | ***None*** | ***Lapse*** |
| CKD | -427.510 | -430.820 | -419.665 | -423.020 | 7.845 | 7.800 |
| ERM | -448.073 | -451.383 | -438.469 | -441.779 | 9.605 | 9.605 |
| KNB | -424.487 | -427.797 | -411.348 | -414.658 | 13.139 | 13.139 |

**Table S1:** Bayes information Criterion (BIC) for fits of Additive and Divisive SDT models to data from **Experiment 3**, with (*Lapse*) and without (*None*) lapse rates.

**TABLE S2**

| **Observer** | **neu** | **con** | **inc** | **$\chi^2$** | ***p*** |
|---|---|---|---|---|---|
| CJD | 0.80 | 0.83 | 0.74 | 4.575 | 0.102 |
| ERM | 0.70 | 0.78 | 0.67 | 5.742 | 0.057 |
| KNB | 0.78 | 0.83 | 0.81 | 1.287 | 0.525 |
| MCO | 0.76 | 0.82 | 0.78 | 2.612 | 0.271 |
| NRB | 0.89 | 0.92 | 0.89 | 1.115 | 0.573 |
| RCL | 0.78 | 0.86 | 0.81 | 3.842 | 0.146 |
| JCO | 0.92 | 0.95 | 0.95 | 1.974 | 0.373 |
| HAP | 0.90 | 0.87 | 0.86 | 1.603 | 0.449 |
| EMW | 0.88 | 0.93 | 0.88 | 3.598 | 0.166 |
| **POOLED** | **0.82** | **0.86** | **0.82** | **15.319** | **<0.001*** ** |

* p < 0.05, ** p < 0.01, *** p < 0.001

**Table S2:** Statistical tests (Pearson's $\chi^2$, *df* = 2) of the hypothesis that the proportion correct (*Pc*) is identical for the *neutral* (**neu**), *congruent* (**con**), and *incongruent* (**inc**) conditions of **Experiment 4**.

**TABLE S3**

| **Observer** | **neu** | **con-0** | $\chi^2$ | ***p*** | **con-180** | $\chi^2$ | ***p*** |
|---|---|---|---|---|---|---|---|
| CJD | 0.80 | 0.90 | 4.800 | 0.028* | 0.75 | 0.982 | 0.322 |
| ERM | 0.70 | 0.83 | 5.905 | 0.015* | 0.72 | 0.129 | 0.720 |
| KNB | 0.78 | 0.84 | 1.500 | 0.221 | 0.81 | 0.362 | 0.548 |
| MCO | 0.76 | 0.83 | 2.185 | 0.139 | 0.81 | 1.15 | 0.283 |
| NRB | 0.89 | 0.96 | 4.579 | 0.032* | 0.87 | 0.142 | 0.706 |
| RCL | 0.78 | 0.86 | 2.736 | 0.098 | 0.85 | 2.068 | 0.150 |
| JCO | 0.92 | 0.97 | 3.241 | 0.072 | 0.92 | 0.022 | 0.883 |
| HAP | 0.90 | 0.88 | 0.280 | 0.597 | 0.86 | 1.061 | 0.303 |
| EMW | 0.88 | 0.98 | 8.422 | 0.004** | 0.88 | 0.000 | 1.000 |
| **POOLED** | **0.82** | **0.89** | **24.383** | **<0.001*** ** | **0.83** | **0.288** | **0.592** |

* $p < 0.05$, ** $p < 0.01$, *** $p < 0.001$

**Table S3:** Statistical comparisons (Pearson's $\chi^2$, *df* = 1) of observer performance in the neutral (**neu**) case of **Experiment 4** with each sub-condition of the congruent trials: aligned-phase (**con-0**) and opposite-phase (**con-180**).

**TABLE S4**

| **Observer** | **con-0** | **con-180** | $\chi^2$ | ***p*** |
|---|---|---|---|---|
| CJD | 0.90 | 0.75 | 7.792 | 0.005** |
| ERM | 0.83 | 0.72 | 3.470 | 0.063 |
| KNB | 0.84 | 0.81 | 0.312 | 0.577 |
| MCO | 0.83 | 0.81 | 0.136 | 0.713 |
| NRB | 0.96 | 0.87 | 5.207 | 0.022* |
| RCL | 0.86 | 0.85 | 0.040 | 0.841 |
| JCO | 0.97 | 0.92 | 2.405 | 0.121 |
| HAP | 0.88 | 0.86 | 0.177 | 0.674 |
| EMW | 0.98 | 0.88 | 7.689 | 0.006** |
| **POOLED** | **0.89** | **0.83** | **15.732** | **<0.001*** ** |

* $p < 0.05$, ** $p < 0.01$, *** $p < 0.001$

**Table S4:** Statistical tests (Pearson's $\chi^2$, $df = 1$) of the hypothesis that proportion correct is identical for phase-aligned (**con-0**) and opposite-phase (**con-180**) stimuli for the congruent case in **Experiment 4**.

### SUPPLEMENTARY FIGURE CAPTIONS

**Figure S1: Effects of density on LTB segmentation thresholds**

Effects of micro-pattern density on segmentation thresholds for experienced observers (author CJD and naïve observers KNB, ERM). Plotted are means (circles) and 1 SEM (lines) obtained from N = 200 bootstrapped fits of the SDT psychometric function. We see that thresholds are slightly higher for 16 micro-patterns, and that similar performance is obtained for 32 and 64 micro-patterns. For each individual observer, 1-way ANOVA on the bootstrapped thresholds revealed a significant effect of density on threshold ($p << 0.001$).

**Figure S2: Thresholds estimated with and without lapse rates**

**(a)** Thresholds from **Experiment 1a**. We see nearly identical threshold estimates whether or not lapse rates are included in our PF definitions. **(b)** Thresholds from **Experiment 1b**.

**Figure S3: Contrast thresholds for LSB segmentation**

(**a**) Fits of SDT psychometric function to LSB segmentation performance for same observers shown in Fig. 3a. (**b**) Histogram of threshold for all observers.

**Figure S4: Thresholds in Experiment 1a and 1b**

Scatterplot of thresholds measured from N = 17 observers segmenting luminance texture boundary (LTB) stimuli using proportion of unbalanced patterns as a cue (**Experiment 1a**), and segmenting luminance step boundary (LSB) stimuli using Michealson contrast (**Experiment 1b**).

**Figure S5: Fits of one-stage model to LSB segmentation performance**

Same as **Fig. 4b** in main text, but shows fits of the one-stage model (**Fig. 4b**) to LSB segmentation data from **Experiment 1b**.

**Figure S6: Fits of additive and divisive one-stage model with lapse**

Same as **Fig. 6c**, but with lapse rates estimated.

**Figure S7: Fits of the divisive model to Experiment 2 data**

Divisive model fit to **Experiment 3** data accurately predicts performance in **Experiment 2**.

**Figure S8: LTB segmentation thresholds for supra-threshold LSB maskers**

(**a**) Three observers segmented LTBs using proportion of unbalanced micro-patterns ($\pi_U$) as a cue in the presence of supra-threshold LSB maskers. LSB masker intensity is plotted in units of the LSB segmentation JND for each observer. For all values greater than 1, the LSB orientation was clearly visible. LTB micro-patterns were presented at both high (0.4, blue curves) and low (0.2, green curves) contrasts. At the higher LTB contrast (0.4), the increase in LTB segmentation thresholds is negligible for observers ERM and KNB, even at the highest masker levels. For observer CJD, the two highest masker levels tested elevated thresholds from about $\pi_U$=0.25 to $\pi_U$ =0.4. On the whole however, we see that LTB segmentation is fairly robust to interference from LSB maskers, consistent with the hypothesis of separate, yet interacting, underlying mechanisms. (**b**) LTBs with various proportions unbalanced micro-patters shown to provide context for interpreting changes in thresholds. Even the largest change we observed, from approximately $\pi_U$ = 0.25 to 0.5, is still fairly subtle.

**Figure. S9: Fits of model in 8a to Experiment 4 data**

Same as **Fig. 8b** in main text, but for remaining observers.

**Fig S10: Analysis of two-stage model results**

(**a**) Bayes Information Criterion, for model selection - favoring two-stage model over the one-stage model ($BIC_2$ - $BIC_1$), for all N = 9 observers in Experiment 4. (**b**) Mean luminance ("DC") response for the first-stage filters obtained from each observer.

**Fig S11: Fits of the two-stage model (Fig. 8a) to Experiment 3 data**

Fits of both additive and divisive instances of the two-stage model shown in **Fig. 8a** to psychophysical data obtained in Experiment 3. As with the one-stage model (**Fig. 6)**, we find much better fits with the divisive two-stage model. Here we were able to simplify the divisive model by eliminating one parameter (setting $\tau_2 = 1$ in **Eq. 4**), and still obtained excellent fits.

**Supplementary Fig. S1**

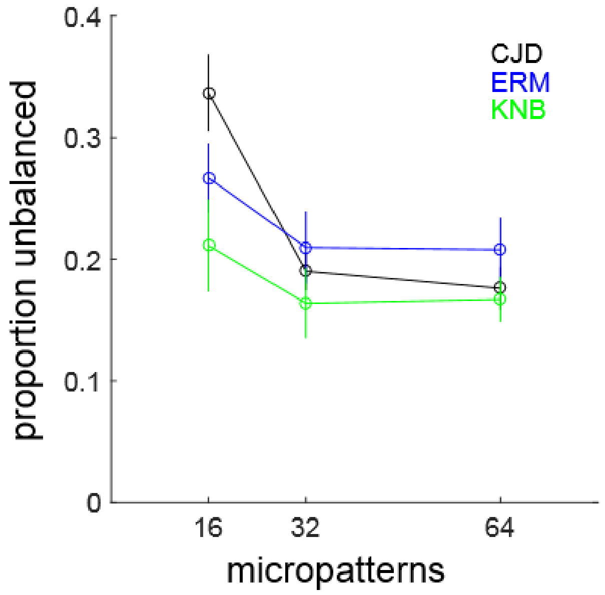

**Supplementary Fig. S2**

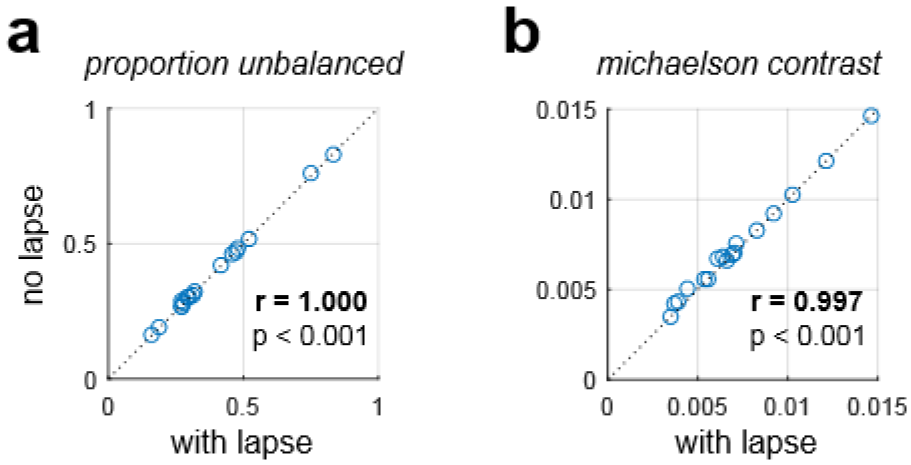

**Supplementary Fig. S3**

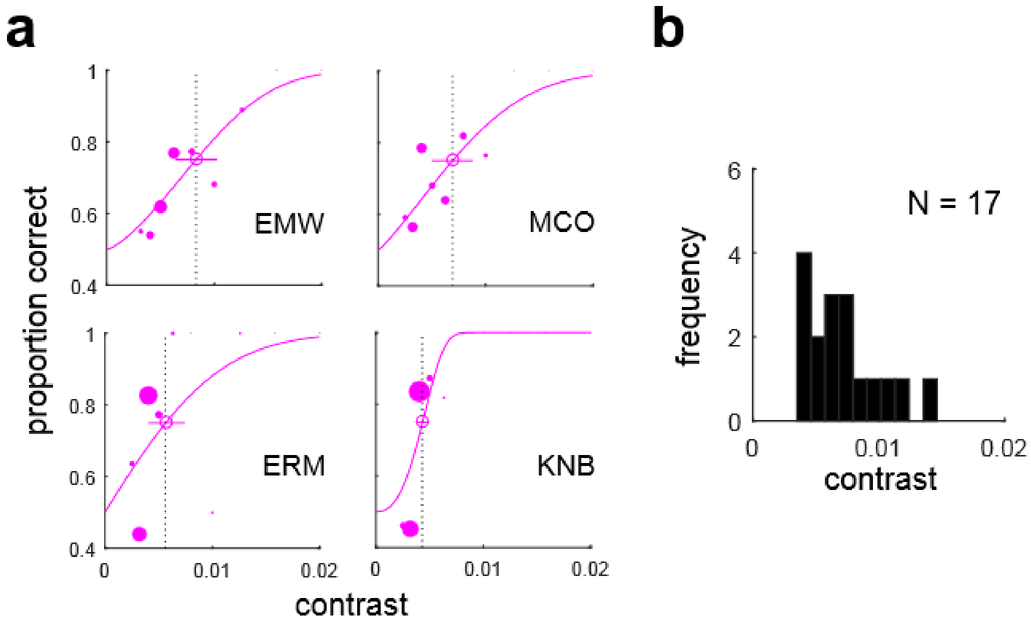

**Supplementary Fig. S4**

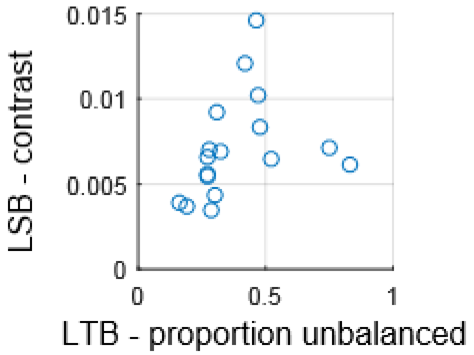

**Supplementary Fig. S5**

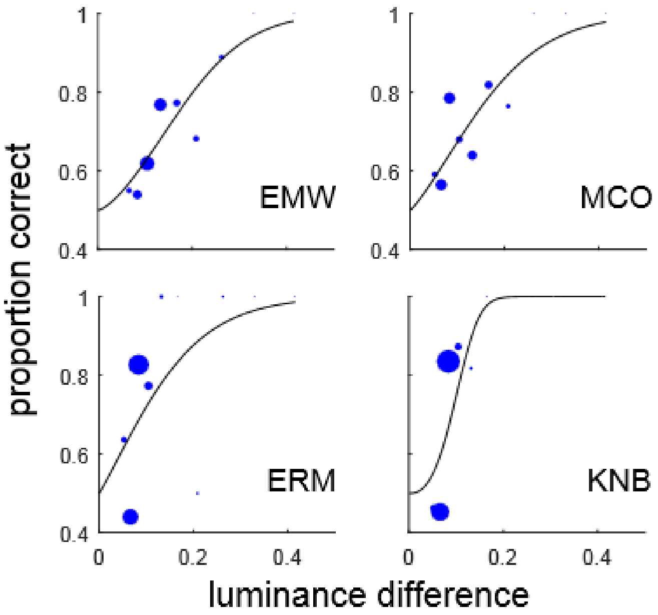

**Supplementary Fig. S6**

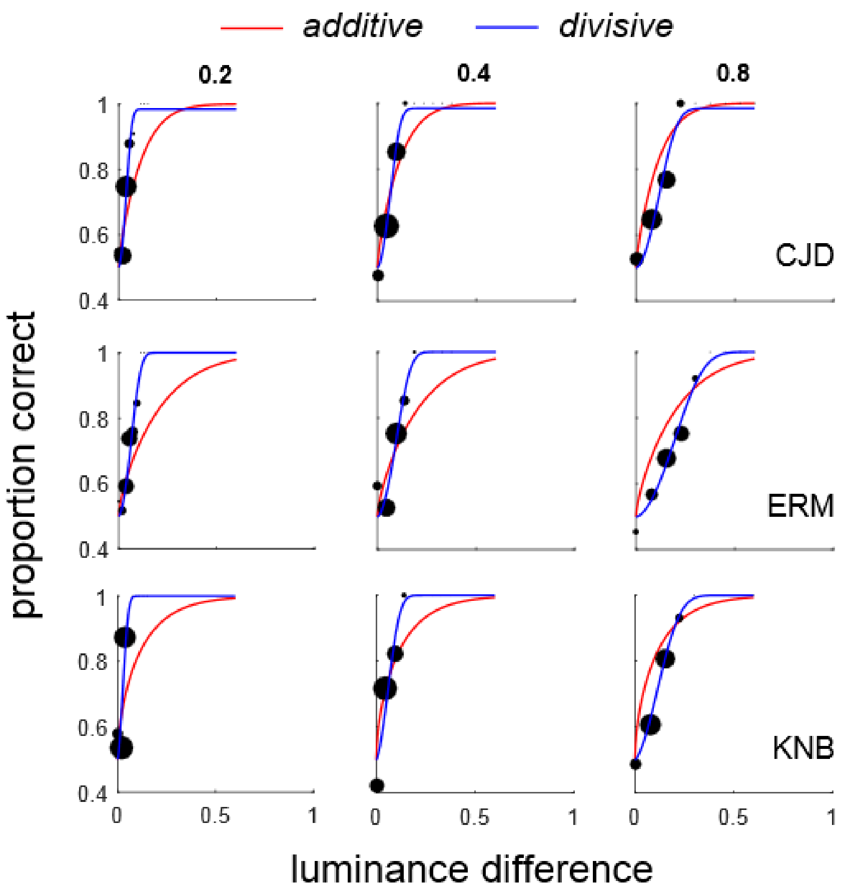

**Supplementary Fig. S7**

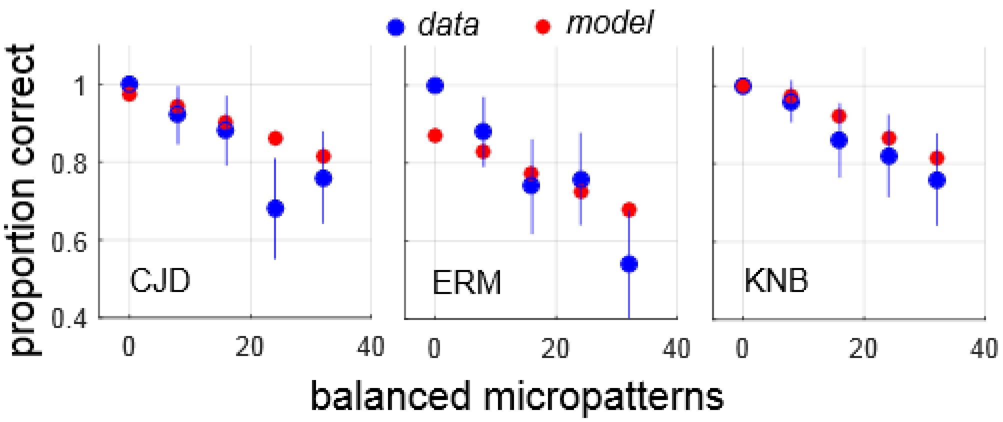

**Supplementary Fig. S8**

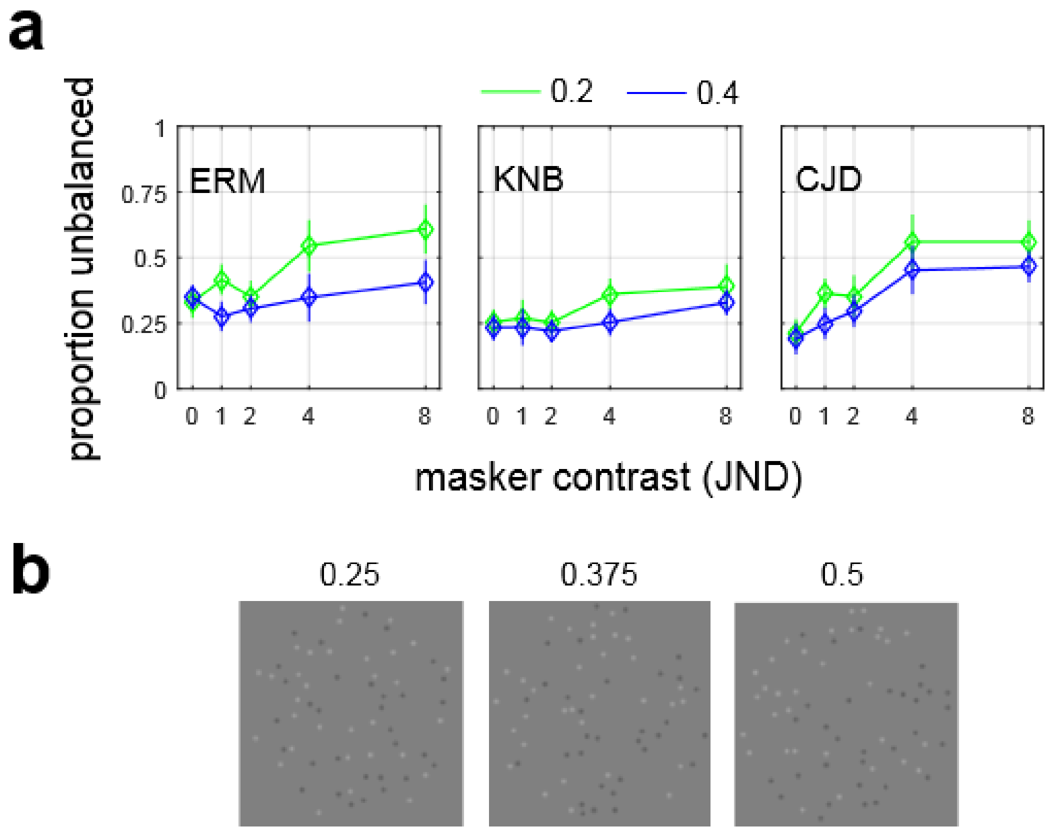

**Supplementary Fig. S9**

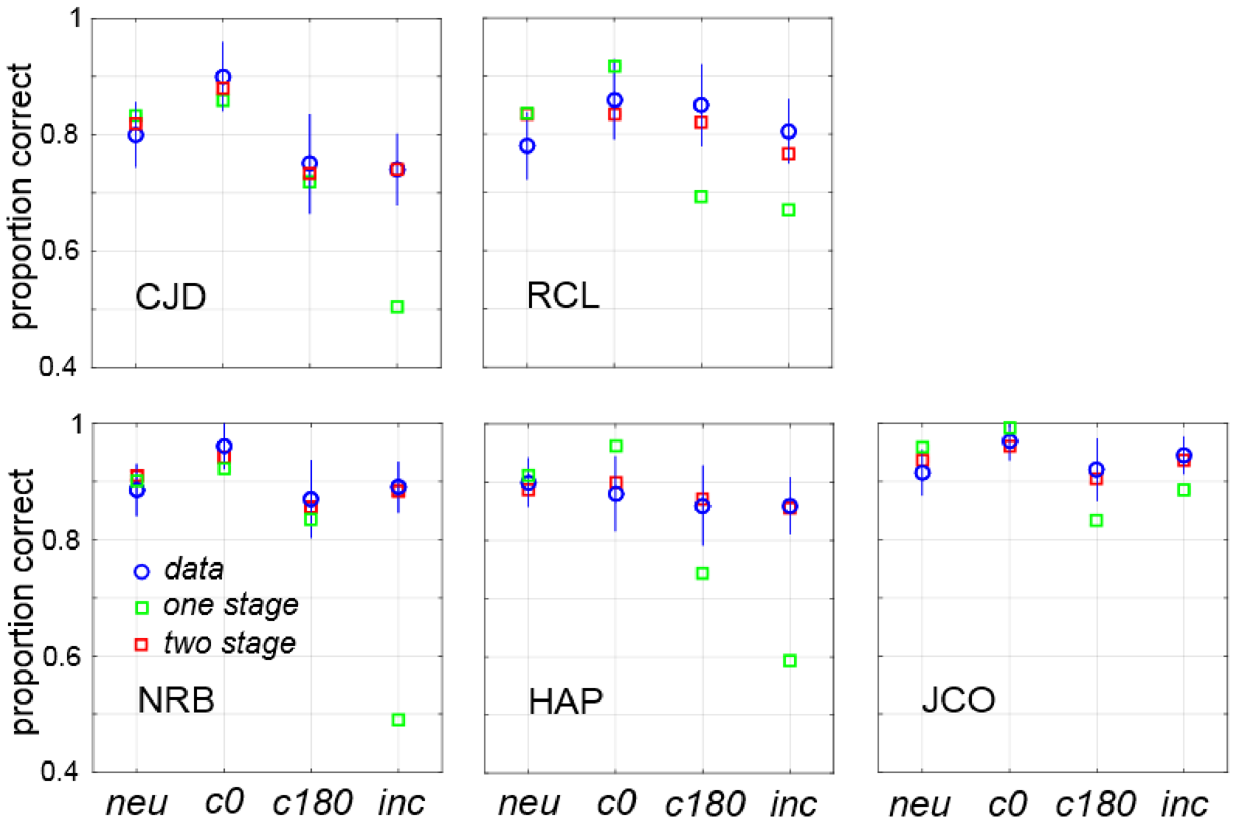

**Supplementary Fig. S10**

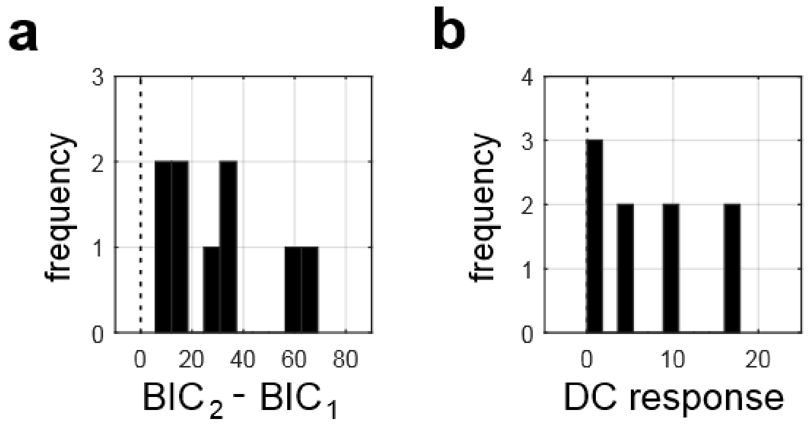

**Supplementary Fig. S11**

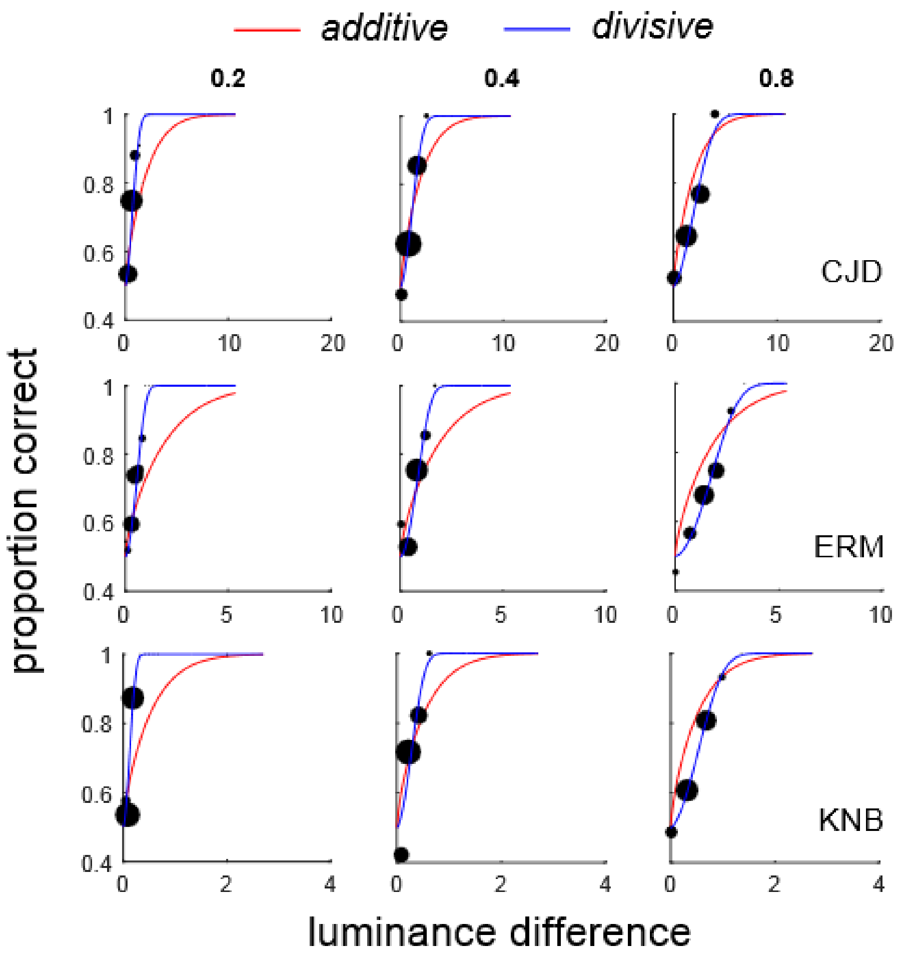
